# Polymicrobial interactions between *Staphylococcus aureus* and *Pseudomonas aeruginosa* promote biofilm formation and persistence in chronic wound infections

**DOI:** 10.1101/2024.11.04.621402

**Authors:** Klara Keim, Mohini Bhattacharya, Heidi A. Crosby, Christian Jenul, Krista Mills, Michael Schurr, Alexander Horswill

## Abstract

Chronic, non-healing wounds are a leading cause of prolonged patient morbidity and mortality due to biofilm-associated, polymicrobial infections. *Staphylococcus aureus* and *Pseudomonas aeruginosa* are the most frequently co-isolated pathogens from chronic wound infections. Competitive interactions between these pathogens contribute to enhanced virulence, persistence, and antimicrobial tolerance. *P. aeruginosa* utilizes the extracellular proteases LasB, LasA, and AprA to degrade *S. aureus* surface structures, disrupt cellular physiology, and induce cell lysis, gaining a competitive advantage during co-infection. *S. aureus* evades *P. aeruginosa* by employing aggregation mechanisms to form biofilms. The cell wall protein SasG is implicated in *S. aureus* biofilm formation by facilitating intercellular aggregation upon cleavage by an extracellular protease. We have previously shown that proteolysis by a host protease can induce aggregation. In this study, we report that *P. aeruginosa* proteases LasA, LasB, and AprA cleave SasG to induce *S. aureus* aggregation. We demonstrate that SasG contributes to *S. aureus* biofilm formation in response to interactions with *P. aeruginosa* proteases by quantifying aggregation, SasG degradation, and proteolytic kinetics. Additionally, we assess the role of SasG in influencing *S. aureus* biofilm architecture during co-infection *in vivo,* chronic wound co-infections. This work provides further knowledge of some of the principal interactions that contribute to *S. aureus* persistence within chronic wounds co-infected with *P. aeruginosa,* and their impact on healing and infection outcomes.

## Introduction

Chronic wound infections contribute to prolonged patient morbidity, with the global burden projected to increase in prevalence over the next decade [1, 2]. It is estimated that chronic wounds such as venous ulcers, pressure ulcers, and surgical site infections impact over 8.2 million people and accrue healthcare costs ranging from $31.7-$96.8 billion in the United States annually [3, 4]. Despite aggressive wound management measures, patients experience treatment failure, wounds that do not heal, and patient morbidity [3, 5, 6]. The primary cause of complications in chronic wounds is the presence of polymicrobial biofilm-associated bacterial infections that lead to prolonged inflammation, collateral tissue damage, and poor vascular perfusion [2, 7–10].

*S. aureus* and *P. aeruginosa* are the pathogens most frequently co-isolated from chronic wound infections, infecting 93.5% and 52.2% of patients, respectively [11–14]. These co-infections are associated with increased bacterial virulence, recalcitrance to treatment, and worsened patient outcomes [15, 16]. Infection severity is exacerbated by competitive interactions that lead to upregulation of exoproducts, surface proteins, and biofilm formation in both pathogens [15, 17–19]. The spatiotemporal dynamics of *S. aureus – P. aeruginosa* co-infections have been well-characterized in chronic infections such as those associated with cystic fibrosis [19–22]. Much remains to be understood about the complex interactions between *S. aureus* and *P. aeruginosa* in the context of chronic wound infections [15].

It has been suggested that these pathogens cannot coexist long-term, and that *P. aeruginosa* ultimately becomes predominant by outcompeting *S. aureus* with its arsenal of anti-staphylococcal exoproducts and higher antimicrobial tolerance [23–26]. However, clinical evidence and recent studies indicate that long-term coexistence between these pathogens occurs frequently, due to coevolution in the wound environment [13, 17, 21]. A predominant ecological theory explaining the infection dynamics between *S. aureus* and *P. aeruginosa* hypothesizes that an initial antagonistic interaction event may occur during early infection, eventually leading to niche partitioning and cooperation [27–29]. Preceding biofilm formation, these two pathogens compete for nutrients and space, leading to antagonism with extracellular products [28, 30]. These interactions with *P. aeruginosa* initiate *S. aureus* biofilm formation independent of host proteins and driven by mechanisms of intercellular aggregation [28]. In response to environmental stress, *S. aureus* often forms free-floating multicellular aggregates, highly tolerant to mechanical disruption and antimicrobial activity [31, 32]. Competitive interactions between *S. aureus* and *P. aeruginosa* serve as a key determinant in establishing chronic infections by enhancing *S. aureus* aggregation and biofilm formation [26, 28]. However, the molecular interactions underlying this response to *P. aeruginosa,* and their impact on chronic wounds has not been clearly defined.

The giant, cell-wall anchored <u>s</u>urf<u>a</u>ce protein <u>G</u> (SasG) has been implicated in *S. aureus* aggregation and biofilm formation [33–35]. SasG has multiple structural domains **(Fig. 1A)** that are orthologous to the *S. epidermidis* accumulation-associated protein (Aap) and function similarly [32, 36–40]. The N-terminal A domain of SasG contributes to adherence by binding to desquamated epithelial cells such as corneocytes [36, 41–43]. The C-terminal B domain of full-length SasG is responsible for aggregation and consists of several B-repeats with alternating G5 subdomains and E spacers [35, 44–47]. We recently demonstrated that the host protease trypsin can induce *S. aureus* SasG-dependent aggregation [48]. Several previous studies indicate that aggregation occurs following cleavage of the A domain by a non-native extracellular protease, which promotes intercellular interactions through Zn^2+^-dependent dimerization of the B repeats [35, 44–47, 49, 50].

**Figure 1.**
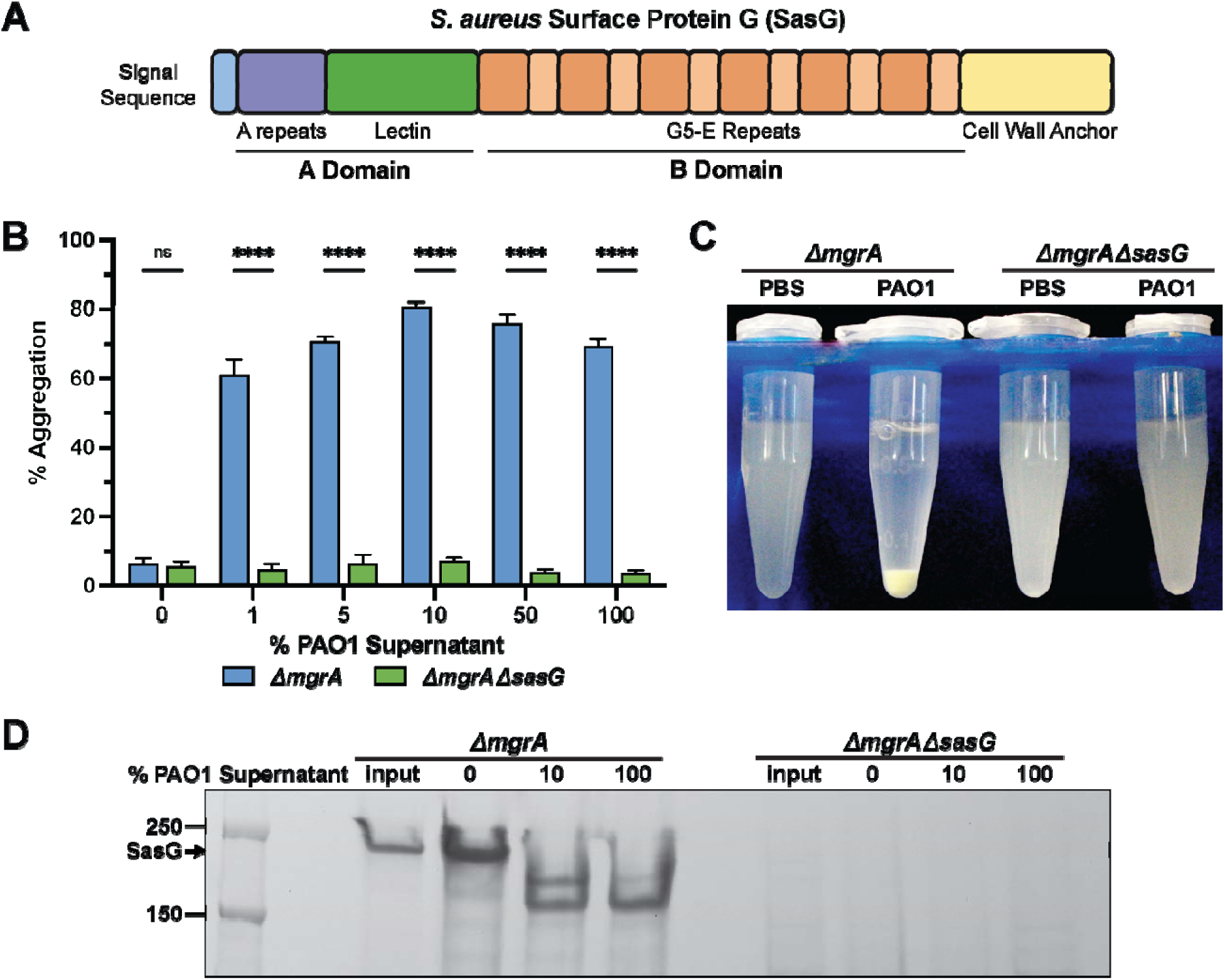
*S. aureus* aggregation is SasG-dependent and induced by *P. aeruginosa*. (A) Graphical representation of the structure of *S. aureus* SasG. (B) Aggregation of *S. aureus* MW2 Δ*mgrA* and Δ*mgrA*Δ*sasG* mutants following incubation for one hour with *P. aeruginosa* cell-free supernatant diluted in PBS in increasing concentrations from 0-100%. (C) Representative image showing aggregation of the Δ*mgrA* and Δ*mgrA*Δ*sasG* mutants following incubation with 10% *P. aeruginosa* supernatant for one hour at room temperature. (D) Coomassie stained SDS-PAGE gel showing SasG processing by increasing concentrations of *P. aeruginosa* supernatant. Cell wall proteins were extracted from the same samples as described above prior to treatment (input) and following aggregation. Results represent an average of three independent experiments performed in triplicate ± SEM (n=9). Statistical significance was determined by 2-way ANOVA with Bonferroni multiple comparisons test (****p<0.0001).

There is variation in SasG expression with most laboratory strains because they either lack functional full-length SasG or do not express it under laboratory conditions [33, 48, 51–53]. However, its clinical relevance is apparent from the identification of anti-SasG antibodies in human serum during infections and of several clinical isolates that express SasG constitutively [51, 54]. Previous work characterizing this mechanism suggests that SasG-dependent aggregation occurs as a protective mechanism to initiate biofilm formation in response to environmental stress [33, 35, 48, 55, 56].

Despite the importance of understanding how antagonistic interactions with *P. aeruginosa* promote *S. aureus* survival and coexistence, little is known about the mechanisms that initiate *S. aureus* biofilm formation during the earliest stages of coinfection. We hypothesize that *P. aeruginosa* proteases cleave SasG and induce *S. aureus* aggregation, which initiates biofilm formation and promotes persistence in polymicrobial chronic wound infections. We propose that SasG-dependent aggregation improves *S. aureus* competitive success in coinfection with *P.* aeruginosa, leading to increases in *S. aureus* survival, antimicrobial resistance, and wound severity. Here we demonstrate that the *P. aeruginosa* proteases LasA, LasB, and AprA cleave SasG and induce *S. aureus* aggregation. We show that SasG-dependent aggregates increase *S. aureus* resistance to antibiotics and promote the formation of robust biofilms that coexist with *P. aeruginosa* in an *in vivo* model of chronic wound infection. These results indicate that SasG plays an important role in the competitive success of *S. aureus* against *P. aeruginosa* and may serve as a crucial mechanism for these pathogens to coexist in chronic infections.

## Results

### Interactions with P. aeruginosa induce SasG-dependent S. aureus aggregation

We recently showed that host proteases can induce *S. aureus* intercellular aggregation by processing the surface protein SasG, conferring protection in chronic lung infections [33, 35]. *P. aeruginosa* secretes several proteases and factors that interact with *S. aureus*, leading us to hypothesize that antagonism by *P. aeruginosa* could induce SasG-dependent aggregation, promoting coexistence in chronic wounds [28, 34, 57, 58]. To investigate this question, we used the previously characterized methicillin-resistant *S. aureus* (MRSA) USA400 MW2 Δ*mgrA* and Δ*mgrA* Δ*sasG* strains [48]. The regulator MgrA represses *sasG* under laboratory growth conditions **(Supplementary Fig. 1A)**; therefore, the Δ*mgrA* mutant was used to evaluate the role of SasG with relevant expression levels [48, 53, 59, 60]. We resuspended the *sasG-* expressing MRSA Δ*mgrA* and the Δ*mgrA* Δ*sasG* double mutant in increasing concentrations (0-100%) of wild-type *P. aeruginosa* PAO1 cell-free supernatant. At all tested concentrations, PAO1 supernatant induced high levels of MRSA Δ*mgrA* aggregation, exhibiting maximum aggregation when treated with 10% supernatant **(Fig. 1B).** Aggregation occurred rapidly, with discernably higher levels of Δ*mgrA* aggregate sedimentation and clearing the suspension within an hour (**Fig. 1C).** As expected, aggregation was abolished in the Δ*sasG* mutant, demonstrating that *P. aeruginosa* induces *S. aureus* aggregation that is dependent on SasG **(Fig. 1B, C, Supplementary Figure 1A)**.

Proteolytic processing within the A domain of SasG is required for *S. aureus* aggregation to occur [35]. We hypothesized that *P. aeruginosa* secreted factors induce *S. aureus* aggregation through processing of SasG. To investigate this we extracted MRSA cell wall proteins following treatment with PAO1 supernatant and evaluated SasG cleavage with SDS-PAGE and Coomassie staining. SasG is anchored to the cell wall at the C-terminally located LPKTG sortase recognition motif [43, 61, 62]. The predicted molecular mass of SasG from strain MW2 is ∼150 kDa, and previous studies observed the protein running to ∼230 kDa, likely due to cell wall remnants from the isolation procedure [32, 39, 59]. Cell wall fragments remain covalently bound to the proteins after extraction due to sortase-anchoring, which slightly impedes migration through the gel [32]. We observed a large protein band at ∼230 kDa in cell wall extracts from Δ*mgrA* not present in Δ*sasG,* which we reasoned to be SasG **(Fig. 1D)**. Treatment with 10-100% PAO1 supernatant also revealed processing into two smaller bands of

∼175 and ∼150 kDa, that were absent in the Δ*sasG* mutant control, indicating processing by PAO1 **(Fig. 1D and Supplementary Fig. 1B).**

To determine if a proteinaceous exoproduct in PAO1 supernatant was responsible for SasG cleavage, we repeated the aggregation assay with heat treated supernatant and observed a loss in Δ*mgrA* aggregation **(Supplementary Fig. 1C).** Since dimerization of exposed B domains facilitates intercellular aggregation, we evaluated the ability of B domain antibodies to inhibit aggregation [35, 38]. Prior to treatment with PAO1 supernatant, we incubated MRSA Δ*mgrA* with antibodies that bind the B domain of SasG and observed inhibition of SasG-dependent aggregation **(Supplementary Fig. 1D)**. These results demonstrate that SasG facilitates *S. aureus* aggregation in response to *P. aeruginosa* secreted factors.

### P. aeruginosa las-regulated proteases induce S. aureus aggregation

*P. aeruginosa* secretes several anti-staphylococcal exoproducts controlled by 3 major quorum-sensing systems, namely *las, rhl* and *pqs* [63, 64]. To determine which *P. aeruginosa* factor(s) process SasG, we generated *las, rhl,* and *pqs* quorum sensing mutants in PAO1. Incubating MRSA Δ*mgrA* with supernatants from PAO1 quorum sensing mutants revealed that Δ*lasR* exhibited significantly attenuated aggregation compared to wild-type PAO1, while Δ*rhlR* and Δ*pqsA* induced high aggregation levels comparable to PAO1 **(Fig. 2A).** Concordantly, recombinant SasG processing assays showed minimal processing by Δ*lasR* supernatant, in contrast to robust processing by PAO1, Δ*rhlR,* and Δ*pqsA* [48] **(Fig. 2B).** Interestingly, though Δ*rhlR* induced significant aggregation, we observed a reduction in SasG processing, likely due to cross-regulation commonly observed between the *las* and *rhl* systems [70, 71] **(Fig. 2C)**. These data indicate *P. aeruginosa* secretes *las*-regulated factors that process SasG to induce *S. aureus* aggregation.

**Figure 2.**
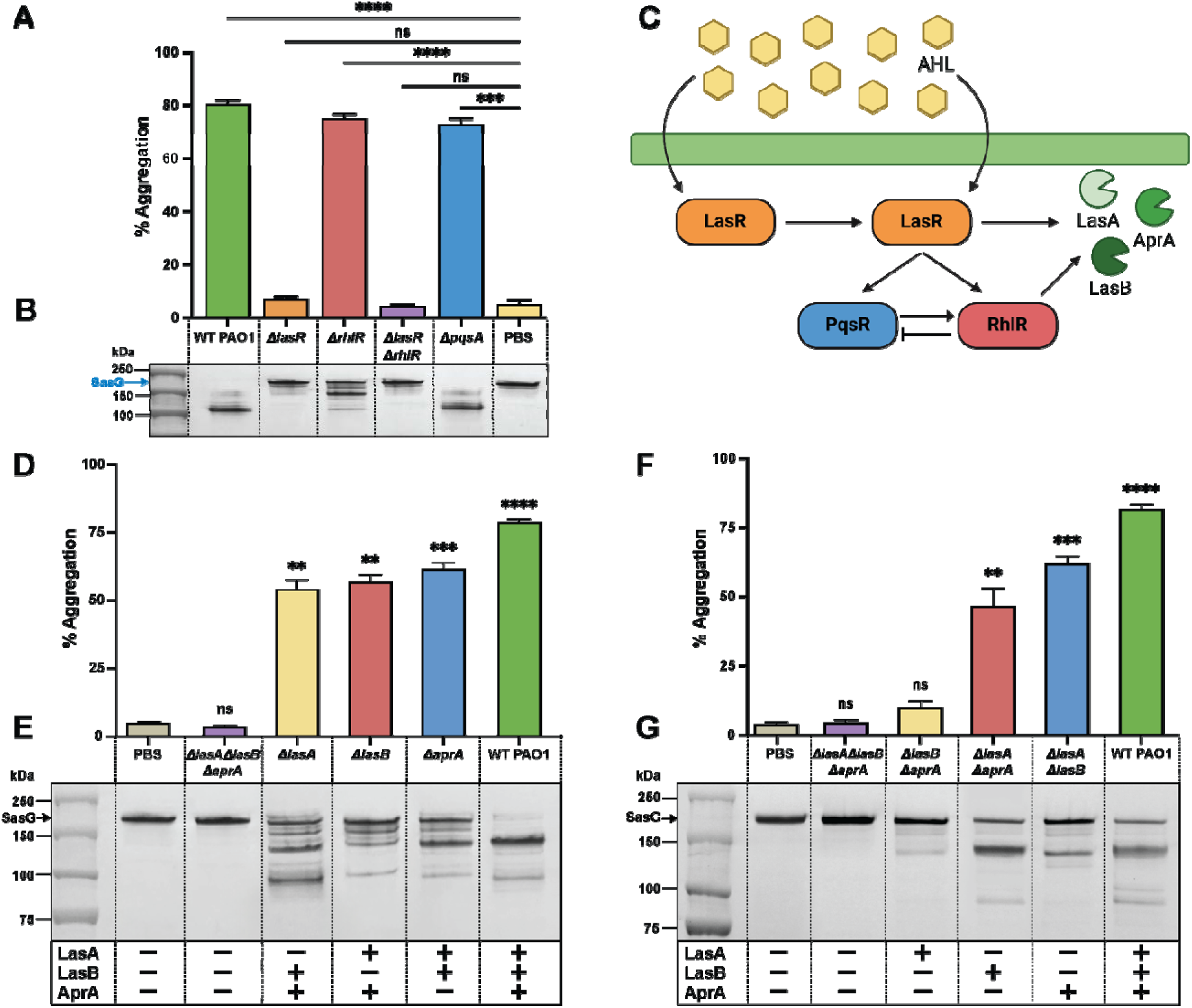
*P. aeruginosa las* regulated proteases cleave SasG and induce *S. aureus* aggregation. (A) Aggregation of the *S. aureus* MW2 Δ*mgrA* mutant treated for one hour with 10% *P. aeruginosa* cell-free supernatant from mutant strains in a PAO1 background with deletions of each of the three major quorum sensing systems (Δ*lasR,* Δ*rhlR,* Δ*pqsA, and* Δ*lasR*Δ*rhlR*). Vertical bars represent four independent experiments SEM ± (n=12). (B) Coomassie stained SDS-PAGE gel showing SasG processing associated with *S. aureus* aggregation induced by PAO1 quorum sensing mutants. (C) MRSA aggregation induced by a triple protease mutant Δ*lasA*Δ*lasB*Δ*aprA*. Each protease is sufficient to cleave SasG, but exhibits varying levels of activity. (D & F) MRSA aggregation induced by various PAO1 protease mutant supernatants. (E & G) SasG cleavage after treatment with 10% PAO1 double protease mutants. Methodology repeated as described in (Fig. 2A & 3B). Vertical bars represent three independent experiments SEM ± (n=9). Statistical significance was determined using the Kruskal-Wallis test with Dunn’s test for multiple comparisons (****p<0.0001, ***p<0.001, **p<0.01, *p<0.1).

### LasA, LasB, and AprA process SasG and induce S. aureus aggregation

The *P. aeruginosa* metalloproteases elastase B (Pseudolysin;LasB), Elastase A (Staphylolysin; LasA), and alkaline protease (Aeruginolysin; AprA) are expressed in a *las* dependent manner and are prolific during early infection, cleaving host and bacterial proteins to facilitate inflammation and clearance of competing bacteria [65, 66]. SasG-dependent aggregation is triggered by proteolytic cleavage by a non-native, extracellular protease, which removes the A domain and exposes the B domain, enabling homodimeric interactions between corresponding B domains on adjacent cell surfaces [34, 35, 48]. To investigate the individual and coordinated contributions of each protease, we performed SasG processing and aggregation assays using supernatants from wild-type PAO1, single protease mutants (Δ*lasA,* Δ*lasB*, and Δ*aprA*), double protease mutants (Δ*lasA* Δ*lasB,* Δ*lasA* Δ*aprA, and* Δ*lasB* Δ*aprA*), and a triple protease mutant (Δ*lasA* Δ*lasB* Δ*aprA*). The triple protease mutant abolished both aggregation (Fig. 2D) and SasG processing (Fig. 2E), demonstrating that at least one of the *P. aeruginosa* proteases, LasA, LasB, or AprA, is responsible for cleaving SasG.

Compared to wild-type PAO1, all single protease mutants (Δ*lasA,* Δ*lasB*, and Δ*aprA*), exhibited attenuated aggregation, indicating that no individual protease alone is sufficient to induce maximal MRSA aggregation **(Fig. 2D)**. SasG processing patterns differed across the double protease mutants, suggesting each protease may target distinct cleavage sites **(Fig. 2E)**. Untreated, recombinant SasG resulted in a ∼165 kDa band, and SasG processing by wild-type PAO1 produced three bands at ∼138 kDa, ∼114 kDa, and ∼100 kDa **(Fig. 2E-G)**. The Δ*lasB* (LasA and AprA) and Δ*aprA* (LasA & LasB) mutants showed slightly reduced processing compared to PAO1, while Δ*lasA* exhibited the most extensive SasG processing, likely due to the combined activity of LasB and AprA **(Fig 2G).** LasB and AprA exhibit functional redundancy and increased coordinated activity when co-expressed, and both proteases exhibit higher expression levels and broader substrate specificities than LasA [72]. Collectively, these data indicate that while LasA, LasB, and AprA each contribute to SasG processing, no single protease alone is sufficient to induce maximal aggregation. Rather, the combined proteolytic activities of LasB and AprA, and to a lesser extent LasA, appear to be primarily responsible for fully processing SasG.

We investigated the ability of each protease to cleave SasG and induce MRSA aggregation using double protease mutants Δ*lasB* Δ*aprA* (LasA^+^), Δ*lasA* Δ*aprA* (LasB^+^), Δ*lasA* Δ*lasB* (AprA^+^). All three proteases were capable of cleaving SasG and inducing MRSA aggregation independently, to varying extents **(Fig. 2F,G).** Both AprA and LasB induced significant aggregation, though moderately attenuatedcompared to wild-type PAO1. In contrast, LasA exhibited limited SasG processing and induced the lowest aggregation levels **(Fig. 2F)**. Despite inducing an intermediate amount of aggregation, Δ*lasA* Δ*aprA* (LasB^+^) processed SasG similarly to wild-type PAO1, producing an intense ∼138 kDa band and a faint ∼100 kDa band **(Fig. 2G)**. Interestingly, Δ*lasA* Δ*lasB* (AprA^+^) induced only slightly less aggregation than PAO1, with reduced processing compared to LasB, exhibiting a less intense ∼138 kDa band, a faint ∼130 kDa band, and several faint bands between 138-165 kDa **(Fig. 2G)**. This increased aggregation by AprA could result from higher activity or cleavage sites more effective at removing the entire A domain than LasB [72]. The Δ*lasB* Δ*aprA* (LasA^+^) induced the lowest aggregation levels and exhibited reduced SasG processing **(Fig. 2G)**. A notable observation was that all double protease mutants produced prominent ∼138 kDa bands when processing SasG, suggesting this may be the location of a primary cleavage site associated with aggregation. We attempted to identify the SasG cleavage site(s) with N-terminal sequencing, with inconclusive results, likely due to the extensive processing resulting in many cleavage sites. Altogether, these findings indicate that while LasA, LasB, and AprA can each independently cleave SasG and induce *S. aureus* aggregation, their combined proteolytic activities likely synergize to fully process SasG, triggering maximal aggregation levels.

### Expression of P. aeruginosa LasA, LasB, and AprA proteases

Previous studies have correlated *P. aeruginosa* protease expression levels with infection severity, finding that quorum-sensing and protease-deficient strains exhibit attenuated virulence in wound models [65, 66, 73, 74]. To investigate which protease(s) are most relevant for polymicrobial interactions and SasG-dependent aggregation in wounds, we quantified *lasA, lasB,* and *aprA* expression in wild-type PAO1 and quorum sensing mutants Δ*lasR*, and Δ*rhlR* using RT-qPCR **(Fig. 3**). Strains were cultured under conditions used for aggregation assays, with transcript levels normalized to the *rpoD* housekeeping gene. In all strains *lasB* was expressed at significantly higher levels than *aprA* and *lasA*, with *lasA* showing the lowest expression levels **(Fig. 3A).** As a major transcriptional activator of these proteases, the *lasR*-deficient mutant exhibited significantly reduced expression of all proteases, which correlated with the previously observed attenuation in SasG processing and MRSA aggregation **Figure 2A-B**. The *rhlR*-deficient mutant also displayed lower protease expression than PAO1, consistent with the reduction in SasG processing observed in **Figure 2B**. LasR was initially identified as the key regulator of protease expression, but the *rhl* quorum sensing system is also required for full activation of some protease genes like *lasB* [71, 75–77]. Therefore, attenuated SasG processing in the Δ*rhlR* mutant is likely a result of reduced expression of *lasB*.

**Figure 3.**
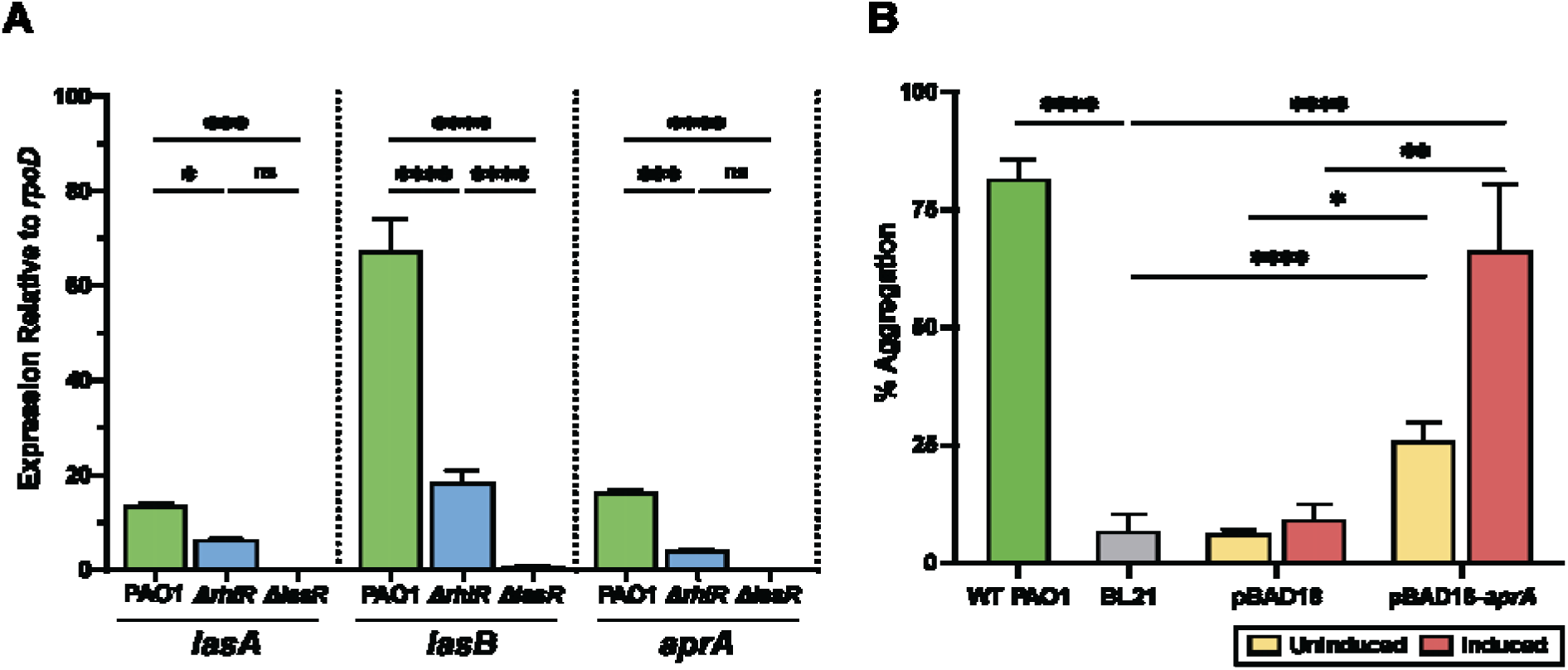
Protease genes *lasA, lasB,* and *aprA* are differentially expressed in *P. aeruginosa* PAO1 in a quorum sensing-dependent manner. (A) Transcriptional expression of protease genes *lasA, lasB,* and *aprA* in wild-type PAO1 and isogenic mutants of Δ*rhlR* and Δ*lasR*. Expression levels were quantified by RT-qPCR, relative mRNA levels for target genes were normalized to the expression of reference gene *rpoD* via the pfaffl method. Vertical bars represent results from 3 independent experiments performed in triplicate SEM ± (n=9). Data were analyzed by 2-way ANOVA with Holm-Sidák multiple comparisons test. (B) AprA induces SasG-dependent aggregation. *E. coli* BL21 expressing the *apr* operon in pBAD18 induced with arabinose or uninduced (repressed with glucose) supernatant was collected and diluted in PBS to 10%. MW2 Δ*mgrA* was treated for 1hr in an aggregation assay and aggregation was quantified. Results represent an average of three independent experiments SEM ± (n=12). Statistical significance was determined using the Kruskal-Wallis test with Dunn’s test for multiple comparisons (****p<0.0001, ***p<0.001, **p<0.01, *p<0.1).

Previous work identified significant upregulation of protease genes, particularly AprA, *in vivo* and in clinical wound specimens, [78]. To validate the relevance of *P. aeruginosa* protease AprA in SasG-dependent aggregation, we expressed AprA in *E. coli* BL21 from an arabinose-inducible promoter. Wild-type BL21 supernatant did not induce MRSA aggregation; however, supernatant from protease over-expressing BL21 induced aggregation to similar levels as wild-type PAO1. These results demonstrate that heterologous expression of AprA is sufficient to induce SasG-dependent aggregation **(Fig. 3B).** Therefore, the proteases LasA, LasB, and AprA are differentially expressed in *P. aeruginosa* and SasG processing is likely the concerted activity of all three proteases, with LasB and AprA being the most prominent in inducing *S. aureus* aggregation.

### Aggregate formation leads to increased S. aureus tolerance to antimicrobials

Chronic wound pathogens experience routine exposure to sub-lethal concentrations of antibiotics, and previous studies indicate that *S. aureus* aggregation promotes antimicrobial tolerance, biofilm formation, and survival post-treatment [28, 31, 57]. Vancomycin and ciprofloxacin are antimicrobials used frequently to treat chronic wounds coinfected with *S. aureus* and *P. aeruginosa* [15, 16]. We investigated if aggregates formed in response to competitive interactions with PAO1 affect *S. aureus* antimicrobial susceptibility and bacterial persistence. The MIC breakpoints against MRSA Δ*mgrA* and Δ*mgrA* Δs*asG* strains for Vancomycin and Ciprofloxacin were 2 µg/mL and 1 µg/mL, respectively. Using the broth microdilution method, MRSA mutant aggregates were exposed to Ciprofloxacin (Cip) **(Figure 4A)** and Vancomycin (Vn) **(Figure 4B)** concentrations ranging from sublethal to 2-4 times the MIC.

**Figure 4.**
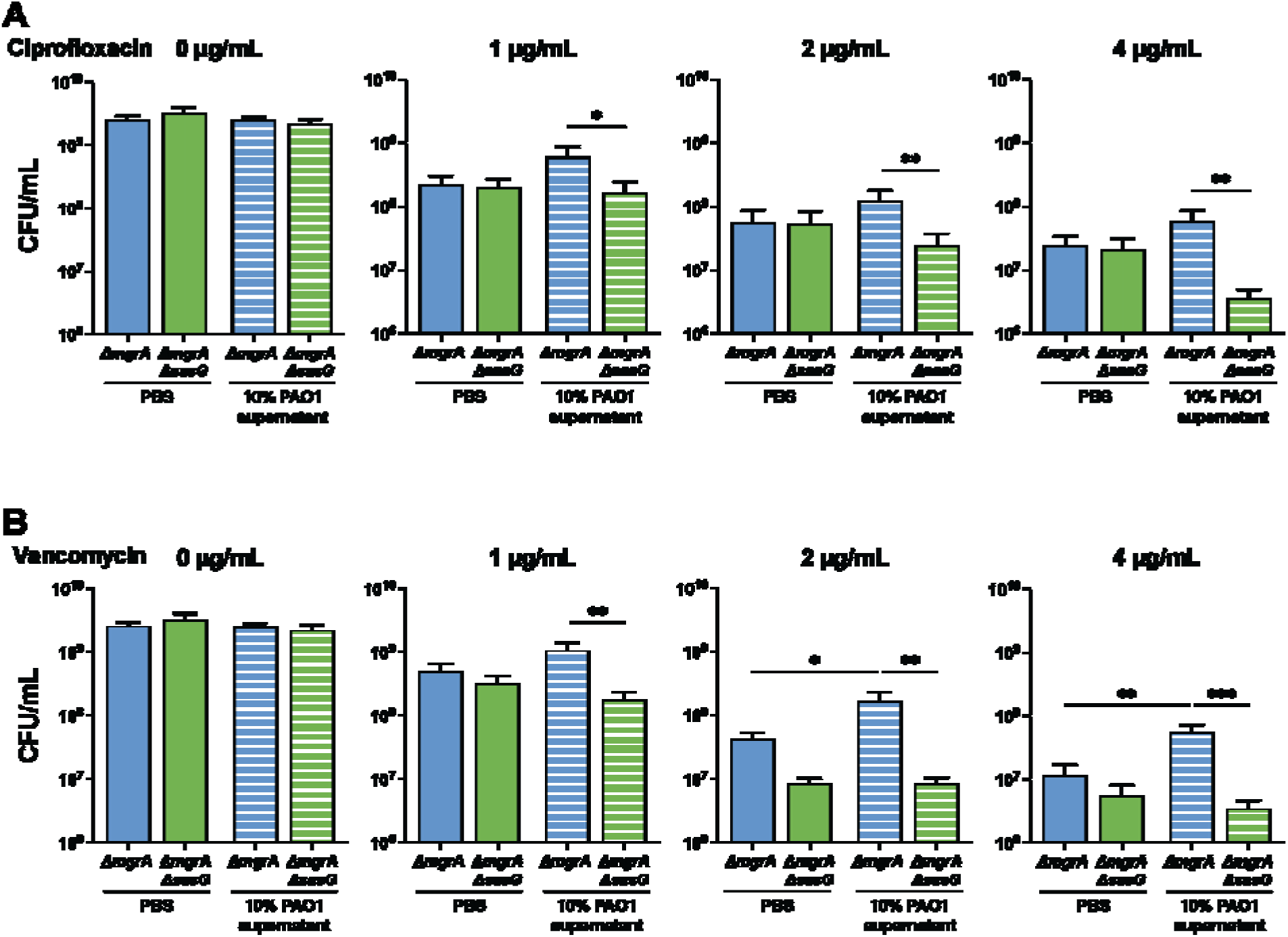
SasG-dependent *S. aureus* aggregates exhibit increased tolerance to antibiotics Ciprofloxacin and Vancomycin. Aggregation assays were performed by treating MRSA ΔmgrA and the Δ*mgrA* Δ*sasG* mutant with either PBS or 10% PAO1 supernatant. Following aggregation for 1hr, MRSA was treated with (A) Ciprofloxacin (Cip) or (B) Vancomycin (Vn) for 5 hours, and CFUs were recovered to quantify antimicrobial susceptibility. Vertical bars represent results from three independent experiments SEM ± (n=9). Statistical significance was determined by One-way ANOVA with Tukey’s multiple comparisons test (****p<0.0001, ***p<0.001, **p<0.01, *p<0.1).

Treatment with PAO1 supernatant facilitated survival of Δ*mgrA* bacteria in a SasG dependent manner, with the Δs*asG* mutant strain exhibiting significantly lower colony forming units (CFUs) compared to Δ*mgrA* control at 2 and 4ug/mL of both antibiotics. Interestingly, the PBS-treated Δ*sasG* mutant also exhibited a significant decrease in cell viability at 2 and 4 µg/mL Vn when compared with Δ*mgrA* bacteria **(Fig. 4B).** We recovered higher CFUs from SasG-expressing MRSA treated with PAO1,than the PBS-treated control, with cell viability at 1 µg/mL Cip and Vn similar to the no antibiotic controls. These results demonstrate that *P. aeruginosa* induced aggregate formation assists the survival of *S. aureus* exposed to ciprofloxacin and vancomycin treatment. **(Fig. 4).**

### S. aureus aggregates promote coexistence during biofilm formation

The earliest stages of coinfection between *S. aureus* and *P. aeruginosa* are crucial in determining if interspecies interactions will lead to coexistence, niche partitioning, or elimination of either pathogen [28, 57]. However, little is known about these interactions, the spatiotemporal dynamics that initiate aggregation and its impact on promoting biofilm formation [22, 28]. We speculated that under the continuous environmental stresses occurring in co-infected wounds, *S. aureus* SasG-dependent aggregates will develop into mature biofilms, capable of coexisting alongside *P. aeruginosa*. The Lubbock Chronic Wound Biofilm Model utilizes wound-like media (WLM) to recapitulate the chronic wound environment *in vitro* [80, 81]. We used this model evaluate the role of SasG in biofilm formation, *S. aureus-P. aeruginosa* interactions, and community spatial organization during early coinfection **(Fig. 5A)**. MRSA Δ*mgrA* and the Δ*sasG* mutants were inoculated into wound-like media (WLM) as either mono-or co-infections with PAO1 and incubated for 24 hours **(Fig. 5A-B)**. We observed no differences in survival among monomicrobial biofilms **(Fig. 5C).** Polymicrobial biofilms consisting of Δ*mgrA* and PAO1 exhibited little difference in cell viability between the two pathogens, and both exhibited slight increases in CFUs compared to the monomicrobial biofilms **(Fig. 5C).** Polymicrobial biofilms with the Δ*sasG* mutant had a significant decrease in MRSA cell viability and an increase in PAO1, suggesting that PAO1 is at an advantage during co-infection with Δ*sasG* **(Fig. 5C)**. In SasG-dependent co-infected biofilms, MRSA made up approximately 50% of the population, which was in sharp contrast to Δ*sasG* biofilms, where MRSA made up less than 5% of the total population **(Supplementary Fig. 3A).**These data suggest that SasG provides *S. aureus* with a survival advantage during coinfection with *P. eruginosa*.

**Figure 5.**
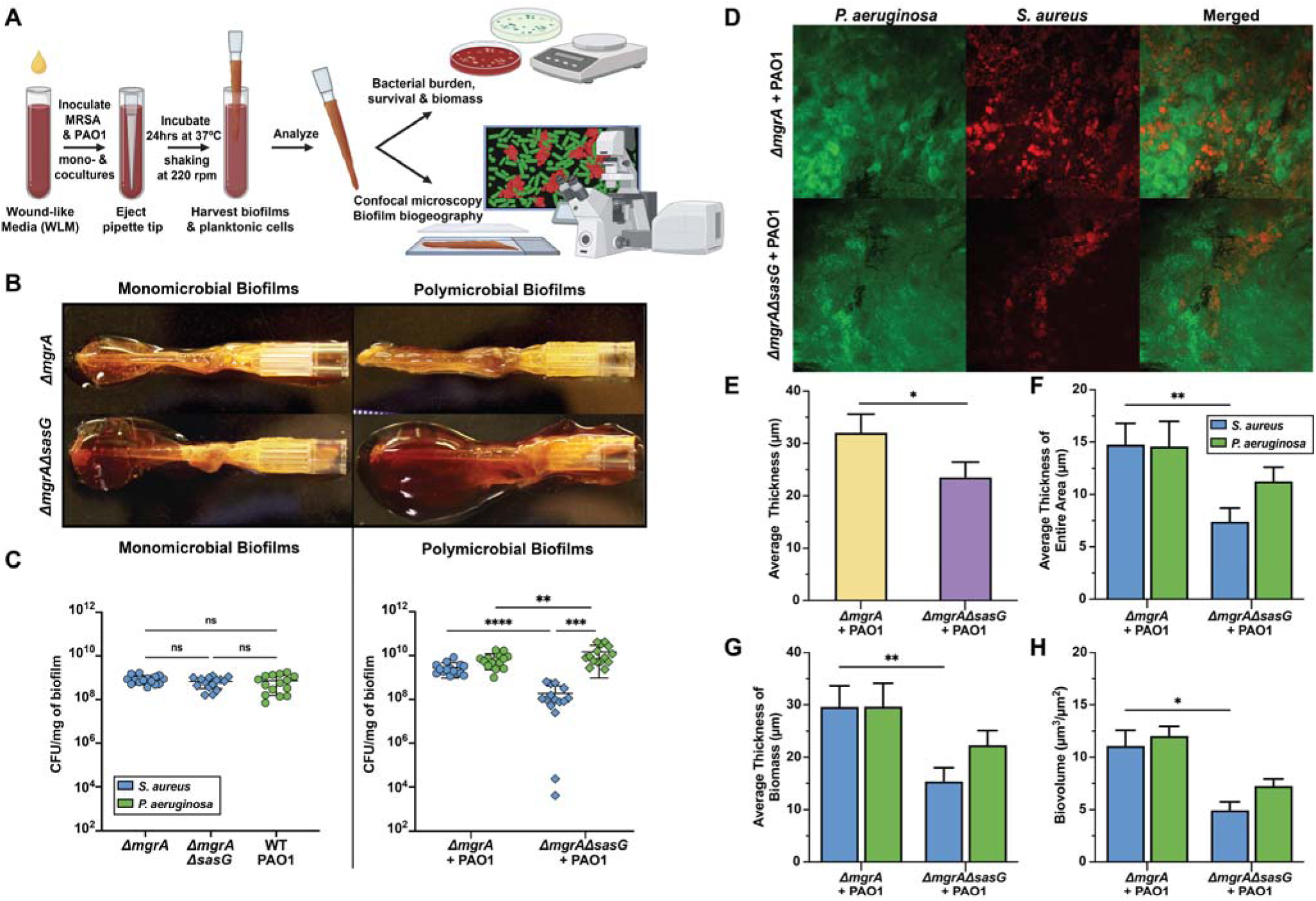
SasG-dependent aggregates promote biofilm formation and contribute to *S. aureus* survival when co-infected with *P. aeruginosa in vitro*. (A) Schematic of the Lubbock Biofilm model. (B) Representative images of mono-microbial and polymicrobial biofilms formed in the Lubbock Model. (C) MRSA and PAO1 CFUs recovered from monomicrobial and polymicrobial biofilms inoculated with MRSA ΔmgrA, the Δ*mgrA* Δ*sasG* mutant, and/or PAO1 represented by CFU/mg of biofilm. (D) Representative confocal microscopy images of polymicrobial biofilms expressing MRSA-dsRED or PAO1-GFP, taken with the 60X objective. (E) Average total thickness of polymicrobial biofilms (µm). (F) Average thickness of the entire biofilm area taken up by MRSA or PAO1 (µm). (G) Average thickness of the MRSA or PAO1 biomass within the biofilm (µm) (H) Total biovolume of MRSA or PAO1 within Lubbock Biofilms (µm^3^). All microscopy images were quantified with COMSTAT and results represent an average of three independent experiments performed in triplicate SEM ± (n=9). Statistical significance of CFU recovery was determined by One-way ANOVA with Tukey’s multiple comparisons test, and image analyses were determined by Mann-Whitney test SEM ± (n=9). (****p<0.0001, ***p<0.001, **p<0.01, *p<0.1).

To investigate how SasG-dependent biofilm formation contributes to spatial structure and *S. aureus* coexistence with *P. aeruginosa*, fluorescent strains of MRSA (expressing pHC48; dsRed) and PAO1 (expressing pMRP9-1; GFP) were inoculated into WLM as described above. Biofilms were harvested and slides were prepared for confocal laser scanning microscopy (CLSM). In Δ*mgrA*-PAO1biofilms, we observed dense, robust aggregates of *S. aureus* throughout the biomass, interspersed with PAO1 **(Fig. 5D).** The average overall thickness of Δ*mgrA* biofilms decreased significantly in a SasG dependent manner **(Fig. 5E),** while average thickness of the entire area **(Fig. 5F),** average biomass thickness **(Fig. 5G)**, and biovolume **(Fig. 5H)** of MRSA vs PAO1 were nearly equivalent **(Fig. 5F-H).** In PAO1 biofilms containing the Δ*sasG* mutant, we had difficulty identifying MRSA in the biofilm and those identifiable were in distinct niches at the periphery of the biofilm separated from PAO1 **(Fig. 5D)**. The Δ*sasG* mutant made up significantly less of the average thickness, area, biomass thickness, and biovolume **(Fig. 5E-H).** Altogether, these findings suggest that SasG promotes formation of a stable and robust MRSA biofilm composed of large aggregates, allowing MRSA to coexist with *P. aeruginosa*.

### In vivo murine model of polymicrobial chronic wound infections

We developed a murine chronic wound model to investigate the impact of SasG-dependent biofilm formation on *S. aureus* survival during co-infection with *P. aeruginosa* **(Fig 6A)**. Mice were wounded with a 6 mm biopsy punch and mono-or co-infected with PAO1 and either MRSA Δ*mgrA* or the Δ*sasG* mutant. Co-infections of PAO1 with Δ*sasG* and the mono-infections of each strain exhibited very little inflammation and pus over the experiment time course (9 days). By day nine, these wounds were only ∼50% the initial wound size, exhibited scabbing, and had little inflammation remaining. **(Fig. 6B,C)**. Mono-infections of Δ*mgrA* exhibited slower wound healing compared to the other mono-infections, but CFU recovery was nearly equivalent among mono-infected groups **(Fig. 6D,E)**. Coinfections with Δ*mgrA* resulted in a significant delay in wound healing, pus and redness around the wound margins, and macroscopically inflamed skin through day seven **(Fig. 6B,C)**. We recovered significantly less MRSA from Δ*sasG* co-infections than Δ*mgrA*, indicating that SasG promotes *S. aureus* survival in polymicrobial chronic wounds **(Fig. 6D)**. Interestingly, PAO1 survival did not change when comparing co-infections, which suggests *S. aureus* and *P. aeruginosa* coexistence **(Fig. 6E)**. Altogether these data indicate that SasG contributes to MRSA persistence and delayed wound closure in wounds co-infections with *P. aeruginosa*.

**Figure 6.**
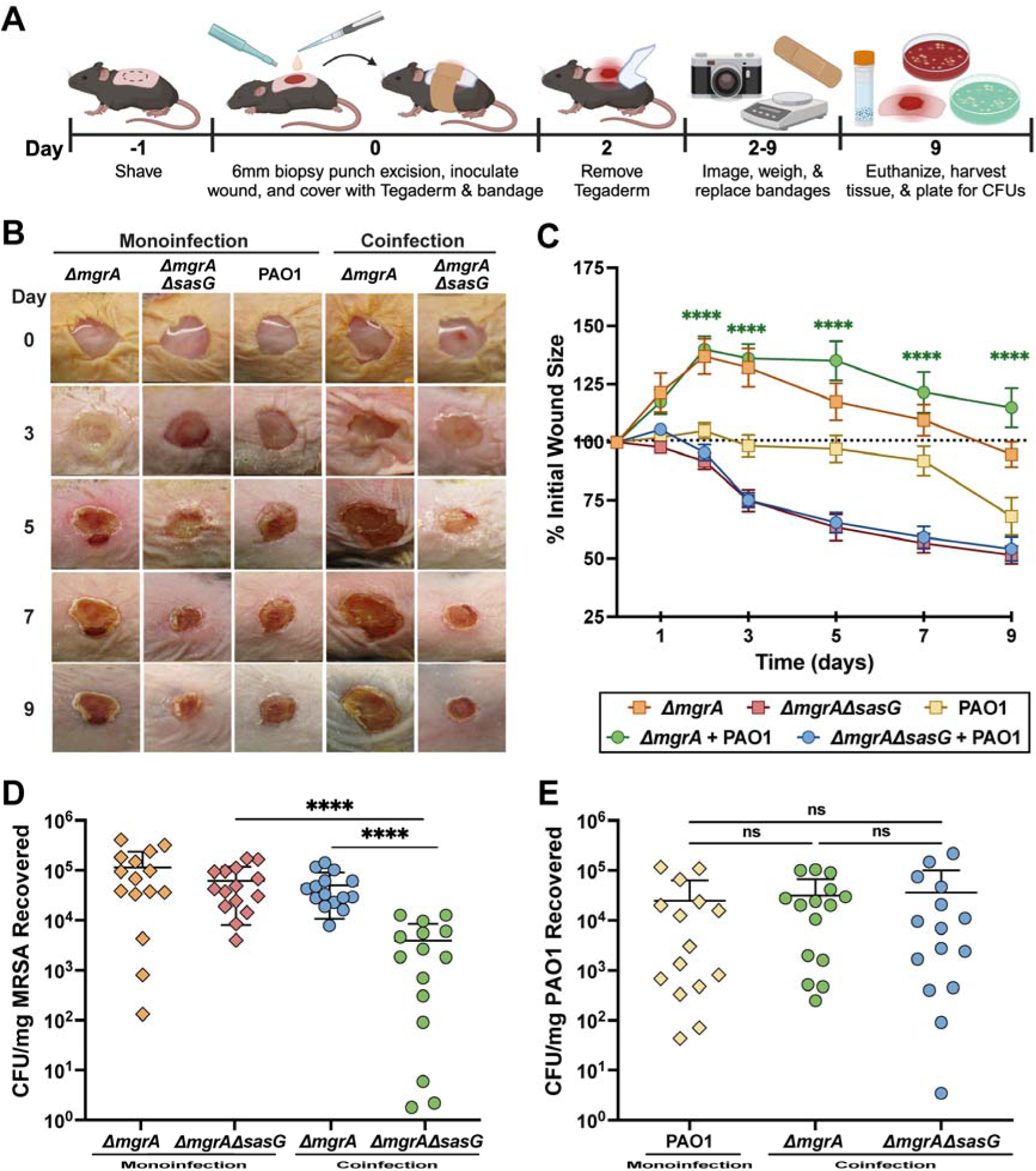
SasG increases *S. aureus* survival and contributes to worse clinical outcomes in an *in vivo* model of polymicrobial chronic wound infections. (A) Schematic of *in vivo* polymicrobial chronic wound model. (B) Representative images showing chronic wound progression over the 9 day time course. (C) Quantification of wound healing over 9 days, represented as the percent difference of the initial wound size with statistical significance representing comparisons between co-infections. (D) MRSA and (E) PAO1 CFUs recovered from excised wound tissue at the day 9 endpoint. Results represent an average of three independent experiments performed SEM ± (n=15). Statistical significance was determined by One-way ANOVA with Tukey’s multiple comparisons test or Mann-Whitney test (****p<0.0001).

## Discussion

Competitive interactions between *S. aureus* and *P. aeruginosa* have been extensively characterized *in vitro* and in chronic lung infections like cystic fibrosis (CF) [19, 22, 30]. However, there is substantial debate and conflicting evidence surrounding their competitive dynamics in chronic wounds [17, 28, 82, 83]. The polymicrobial nature of chronic wounds is well documented in clinical studies, showing *S. aureus* and *P. aeruginosa* co-isolated from wound specimens at a high frequency [84, 85]. This led to the widely accepted view that *S. aureus* promotes secondary *P. aeruginosa* infection but is ultimately outcompeted and displaced, contending that the two species cannot stably coexist [20, 23]. Since the lung of a CF patient is distinct from a chronic wound environment, recent development of novel disease models, both *in vitro* and *in vivo,* has led to work that uncovers the mechanisms of biofilm formation, environmental conditions, and polymicrobial interactions in chronic wounds [17, 21, 86, 87]. These studies provide further evidence that *S. aureus* and *P. aeruginosa* can coexist and describe one mechanism that significantly contributes to this, promoting antimicrobial tolerance, bacterial persistence, and delayed wound healing [17, 21, 86, 87]. Our results show that *P. aeruginosa* may promote *S. aureus* coexistence in chronic wounds by inducing intercellular aggregation upon initial co-infection, encouraging subsequent *S. aureus* biofilm formation.

*P. aeruginosa* upregulates several extracellular factors and proteases in the presence of *S. aureus* that exacerbate tissue damage and delay wound healing [65, 73]. The proteases LasB, AprA, and LasA are found in clinical wound fluid and contribute to delayed wound healing by exacerbating tissue damage, promoting fibrin clot formation, delaying skin restructuring, and encouraging a polymicrobial environment [88]. Our previous work demonstrates that host proteases like trypsin cleave SasG and induce *S. aureus* intercellular aggregation [48, 59]. This led us to hypothesize that SasG-dependent intercellular aggregation serves as a protective mechanism against polymicrobial interactions, facilitating swift *S. aureus* biofilm formation in response to antagonism by *P. aeruginosa*. Here, we demonstrate that secreted proteases in *P. aeruginosa* supernatant cleave SasG and induce aggregation **(Fig. 1B-D).** This aggregation was attenuated by heat-treating the supernatant, validating the involvement of proteinaceous factors **(Supplementary Fig. 1B).**

LasR is generally considered the master regulator of LasA, LasB, and AprA [63, 64, 89]. As expected, in **Figure 2** Δ*lasR* eliminated aggregation and SasG processing, but we also observed a slight reduction or delay in SasG processing by Δ*rhlR.* This is likely explained by the interconnected nature of the *rhl* and *las* quorum sensing systems. Previous work found that *rhl* compensates for virulence factor expression if the *las* quorum sensing system is disrupted [71, 76]. RhlR is also required for full activation of some virulence factors, so the attenuated SasG processing **(Fig. 2B)** and low gene expression **(Fig. 3A)** is likely due to incomplete activation of *lasB* or other genes reliant on *rhl* [70, 90–92].

Our previous work identified a metalloprotease, SepA, in *S. epidermidis* that cleaves the SasG ortholog Aap and induces aggregation following a similar mechanism [39]. *P. aeruginosa* metalloproteases LasA, LasB, and AprA are detected at high concentrations in chronic wound infections breaking down host matrix molecules, which further develops an environment ideal for polymicrobial interactions [39, 48], and we confirmed that no significant *S. aureus* aggregation or SasG processing occurred when exposed to the triple protease mutant (Δ*lasA*Δ*lasB*Δ*aprA*) supernatant.

Evaluation of single protease mutant phenotypes highlighted the synergy between proteases. The attenuated aggregation and extensive SasG processing observed in Δ*lasA* could result from SasG overprocessing by LasB and AprA **(Fig. 2D-E).** LasB and AprA can act in concert to enhance proteolytic activity, which may also explain the enhanced SasG cleavage by Δl*asA* [93]. Double protease mutants exhibited greater variations in SasG cleavage and aggregation than single mutants **(Fig. 2F-G).** LasA (Δ*lasB* Δ*aprA*) exhibited the weakest activity, likely due to its limited specificity and need for activation by LasB [94]. LasA has a narrow substrate specificity and cleaves glycine-rich substrates, preferring bonds in Gly-Gly-Ala sequences [95, 96]. The SasG A domain sequence contains only a single predicted cleavage site with these residues, so LasA may ineffectively remove the A domain compared to LasB and AprA. SasG processing by double protease mutants resulted in multiple cleavage events and several bands with varying molecular weights **(Fig. 2E & G)**. We attempted to identify proteolytic cleavage sites using N-terminal sequencing, but results were inconclusive, even in shorter reactions with only one presumable cleavage event (data not shown). SasG cleavage was not limited to a single defined product as seen previously with human trypsin and *S. epidermidis* SepA [39, 48]. Based on the structure of SasG and previously identified cleavage sites, LasA, LasB, and AprA likely cleave at multiple sites within the lectin portion of the A domain to allow B domain dimerization [39, 97]. Overall, our data indicate that the combined activities of LasA, LasB, and AprA are required for maximal SasG processing and aggregation.

Aggregation and competitive interactions between *S. aureus* and *P. aeruginosa* can promote synergism and alter antimicrobial tolerance [17, 19, 98]. Bacterial aggregates often exhibit characteristics similar to mature biofilms, such as altered metabolism, gene expression, and protection from environmental stress [57]. Our data demonstrate that SasG-dependent aggregation induced by PAO1 increased MRSA tolerance to vancomycin and ciprofloxacin, which are commonly used to treat *S. aureus-P. aeruginosa* coinfections [17]. The increased antimicrobial tolerance of Δ*mgrA* is likely attributable to the protective effects conferred by aggregate formation, preventing effective antibiotic interaction with the cell surface. Previous work demonstrated that *P. aeruginosa* factors can synergize with or antagonize antibiotic activity against *S. aureus* in a strain-dependent manner [19, 22, 99, 100]. LasA was reported to protect *S. aureus* from vancomycin *in vivo* while potentiating killing *in vitro* [19, 98]. Therefore, the enhanced antimicrobial tolerance observed for Δ*mgrA* likely results from a combination of aggregate formation plus *P. aeruginosa* proteases and secreted factors reducing antibiotic efficacy. Conversely, the increased susceptibility of the Δ*sasG* mutant could stem from *P. aeruginosa* factors potentiating antimicrobial effects in the absence of SasG-mediated aggregation. The interplay between SasG expression and *P. aeruginosa* interactions may provide *S. aureus* with a competitive advantage by conferring protection from environmental stresses and enabling stable co-existence within the chronic wound environment.

Bacterial aggregates often provide increased stability and protection compared to polysaccharide biofilms, but their contribution to long-term biofilm development and community organization in chronic wounds remains poorly understood [55]. Using the Lubbock Chronic Wound Biofilm model, we demonstrated that SasG contributed to *S. aureus* biofilm formation and long-term survival in co-infections with *P. aeruginosa.* Previous work investigating biofilm biogeography in wounds observed patchy distributions of each bacterial population, with the majority of *S. aureus* biomass identified as aggregates, driving *P. aeruginosa* into planktonic cells [101]. We observed a similar community structure, and confocal microscopy revealed MRSA Δ*mgrA* forming biofilms made up of dense SasG-dependent aggregates interspersed among populations of *P. aeruginosa* **(Fig. 5D).** We observed several dual species aggregates and found that *S. aureus* aggregates grew in close proximity to or within *P. aeruginosa* populations, suggesting stable coexistence between the pathogens. Previous studies observed niche partitioning of *S. aureus* and *P. aeruginosa* during coinfection, which we observed with Δ*sasG* but *P. aeruginosa* dominated in these biofilms [22], Interestingly, the Δ*sasG* biofilms appeared as a large blood clot compared to the smaller, dense Δ*mgrA* polymicrobial biofilms **(Fig. 5B).** We speculate that *S. aureus* Δ*sasG* mutant forms biofilms through coagulation and clumping mechanisms as opposed to intercellular aggregates [57, 102].

Biofilm formation functions to protect bacteria from host immune factors, antimicrobial molecules, and competitors, which is a crucial component to persistence and treatment failure in chronic wound infections. Previous clinical and *in vivo* studies of chronic infections demonstrate increased virulence, biofilm formation, and persistence in coinfections of *P. aeruginosa* with *S. aureus* [17]. Using an *in vivo* model of chronic wound infections we observed a significant delay in wounds coinfected with *P. aeruginosa* and *S. aureus* Δ*mgrA* **(Fig. 6B-C).** Interestingly, we also observed a slight delay in wound healing in Δ*mgrA* mono-infections, indicating additional potential SasG cleavage by host proteases, such as matrix metalloproteases.. We observed significant attenuation of *S. aureus* survival in Δ*sasG*-PAO1 coinfections, while Δ*mgrA* coinfections exhibited equivalent *P. aeruginosa* and *S. aureus* populations. We speculate that upon coinfection, close contact between the pathogens allows competitive interactions to occur and initiates SasG-dependent aggregation. Our findings indicate that *S. aureus* establishes SasG-dependent biofilms that allow for persistence. Ultimately *S. aureus* SasG-dependent biofilm formation and coexistence with *P. aeruginosa* result in recalcitrant chronic wound infections that exhibit delayed wound healing, reduced antimicrobial efficacy and poor patient outcomes. Altogether our findings demonstrate a novel mechanism for how *P. aeruginosa* facilitates co-existence wtih *S. aureus* in wounds specifically through the activity of proteases LasA, LasB, and AprA which induce SasG-dependent aggregation.

## Materials and Methods

### Ethics Statement

All animal studies described were reviewed, approved, and done in accordance with the recommendations of the Animal Care and Use Committee at the University of Colorado Anschutz Medical Campus. The approved protocol was assigned number 00987.

### Bacterial Strains, Media, and Growth Conditions

All bacterial strains and plasmids used in this work are listed in **Table 1**. In this work we used mutant strains of *S. aureus* MRSA USA400 MW2 as described in our previous work[48], that were either SasG-expressing (MRSA Δ*mgrA*) or a SasG mutant (MRSA Δ*mgrA*Δ*sasG*). The regulator MgrA represses *sasG* under laboratory conditions, so this mutant is used to better reflect clinically relevant expression levels in our investigation under laboratory conditions [48, 53, 59, 60]. *S. aureus* strains were inoculated into tryptic soy broth (TSB; BD), and strains of *E. coli* and *P. aeruginosa* were inoculated into Lennox lysogeny broth (LB; RPI) unless indicated otherwise. Cultures were grown overnight at 37°C with shaking at 200 rpm. Antibiotics were added to the media at the following concentrations: chloramphenicol (Cam) 10 µg/mL and erythromycin (Erm) 5 µg/mL. *E. coli* strains with plasmids were maintained in LB supplemented with antibiotics at the following concentrations: ampicillin (Amp) 100 µg/mL, gentamicin (Gn) 20 µg/mL, and chloramphenicol (Cam) 10µg/mL.

**Table 1.**
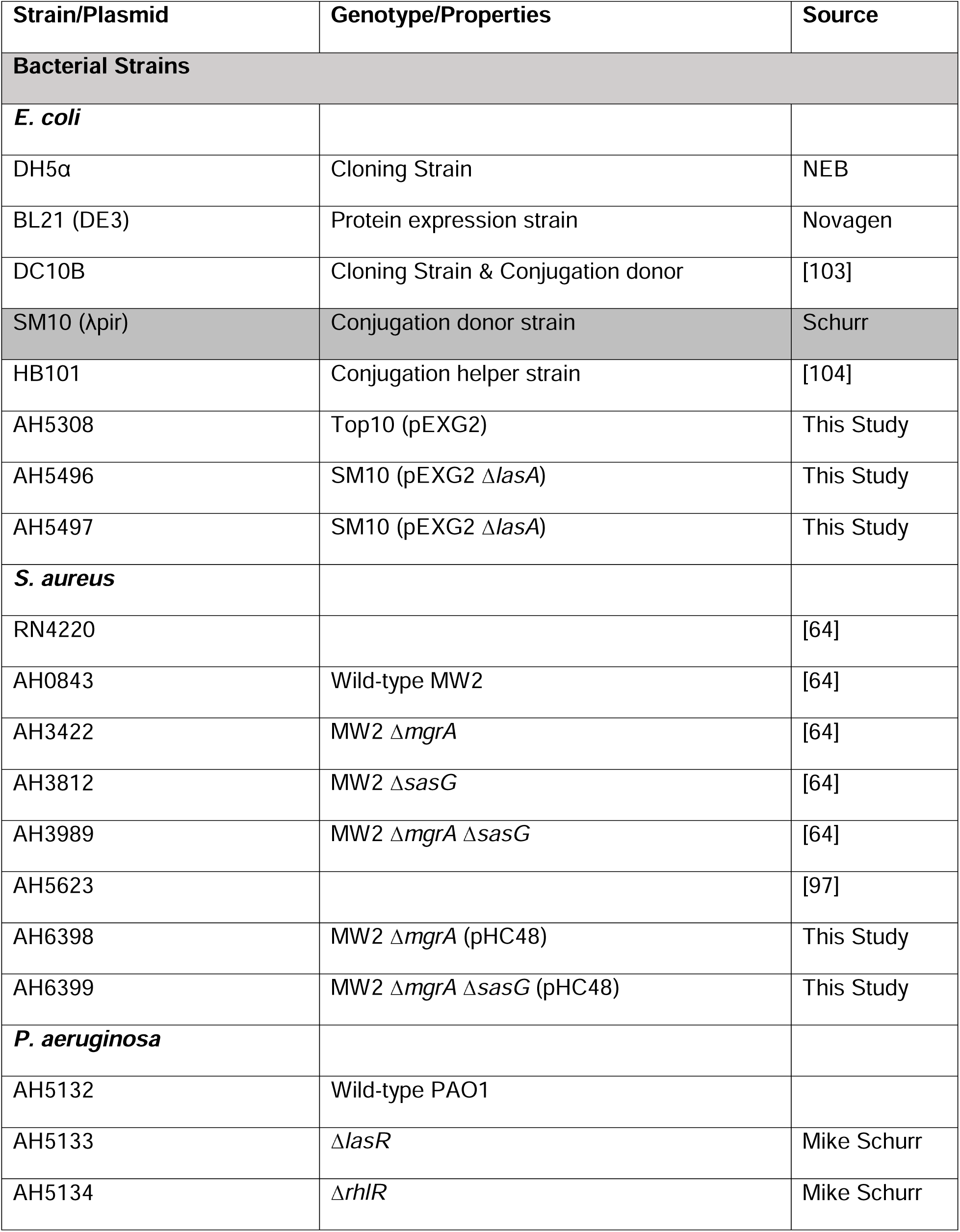
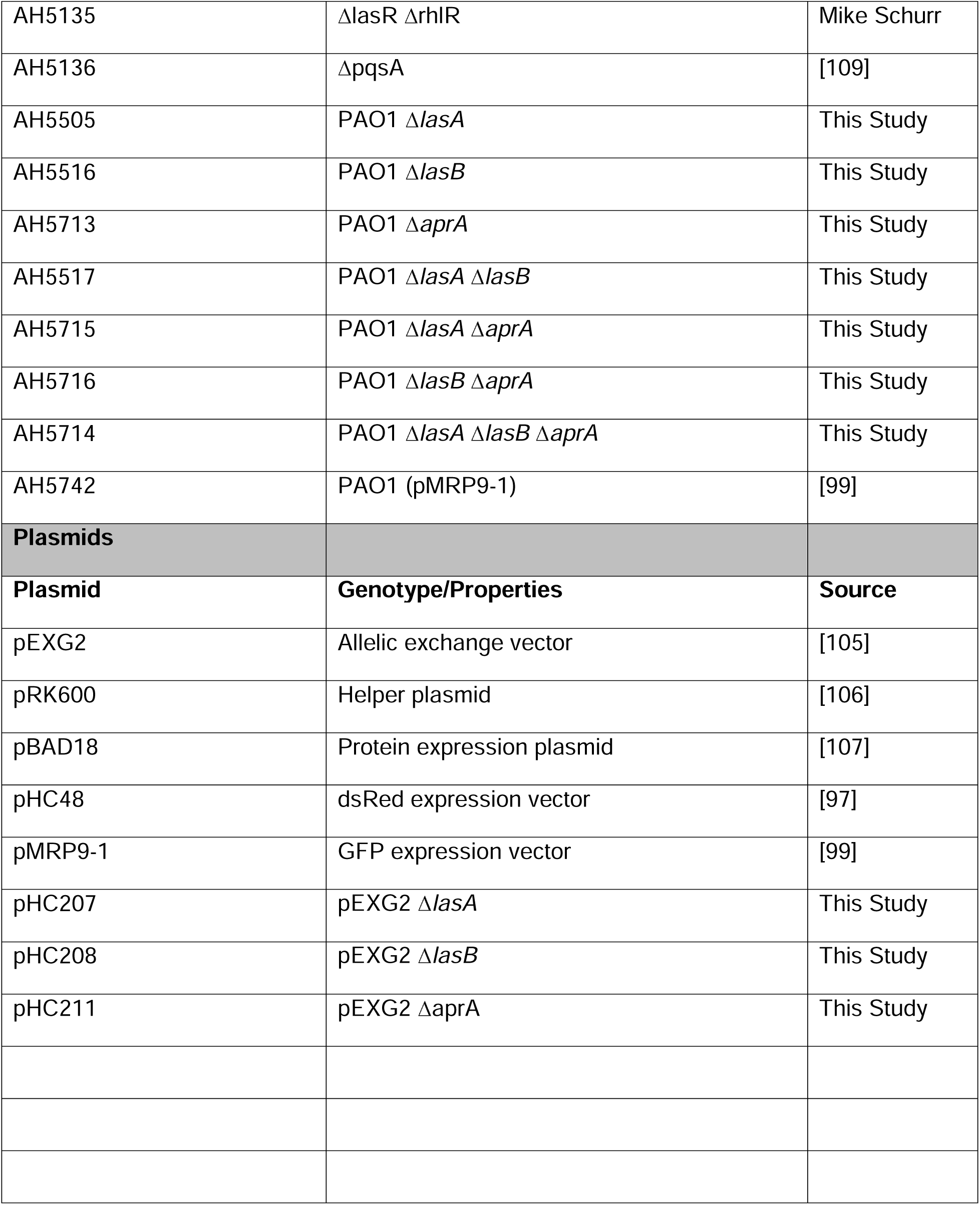
Bacterial strains & plasmids.

### Recombinant DNA and Genetic Techniques

*E. coli* DH5α and DC10B were used as cloning hosts for plasmid construction. All restriction enzymes, Phusion and Q5 polymerases, and DNA ligase were purchased from New England Biolabs (NEB). Kits for DNA extraction, plasmid mini-preps, and gel extractions were purchased from Qiagen. Lysostaphin and lysozyme were purchased from Sigma-Aldrich and used for DNA extractions. All oligonucleotides were purchased from Integrated DNA Technologies (IDT) and are listed in **Table 2**. DNA sequencing was performed at the Molecular Biology Service Center at the University of Colorado Anschutz Medical Campus.

**Table 2.**
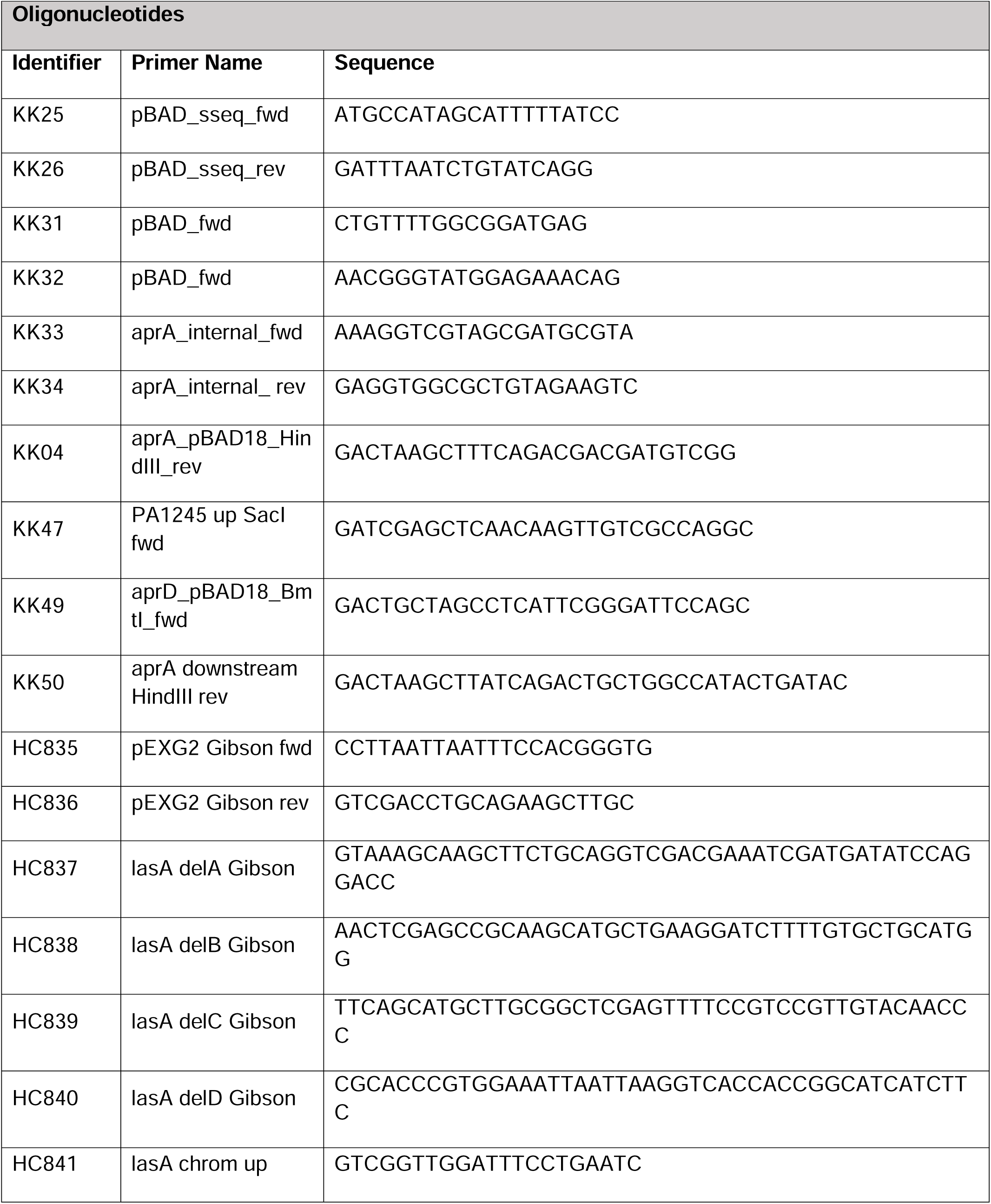
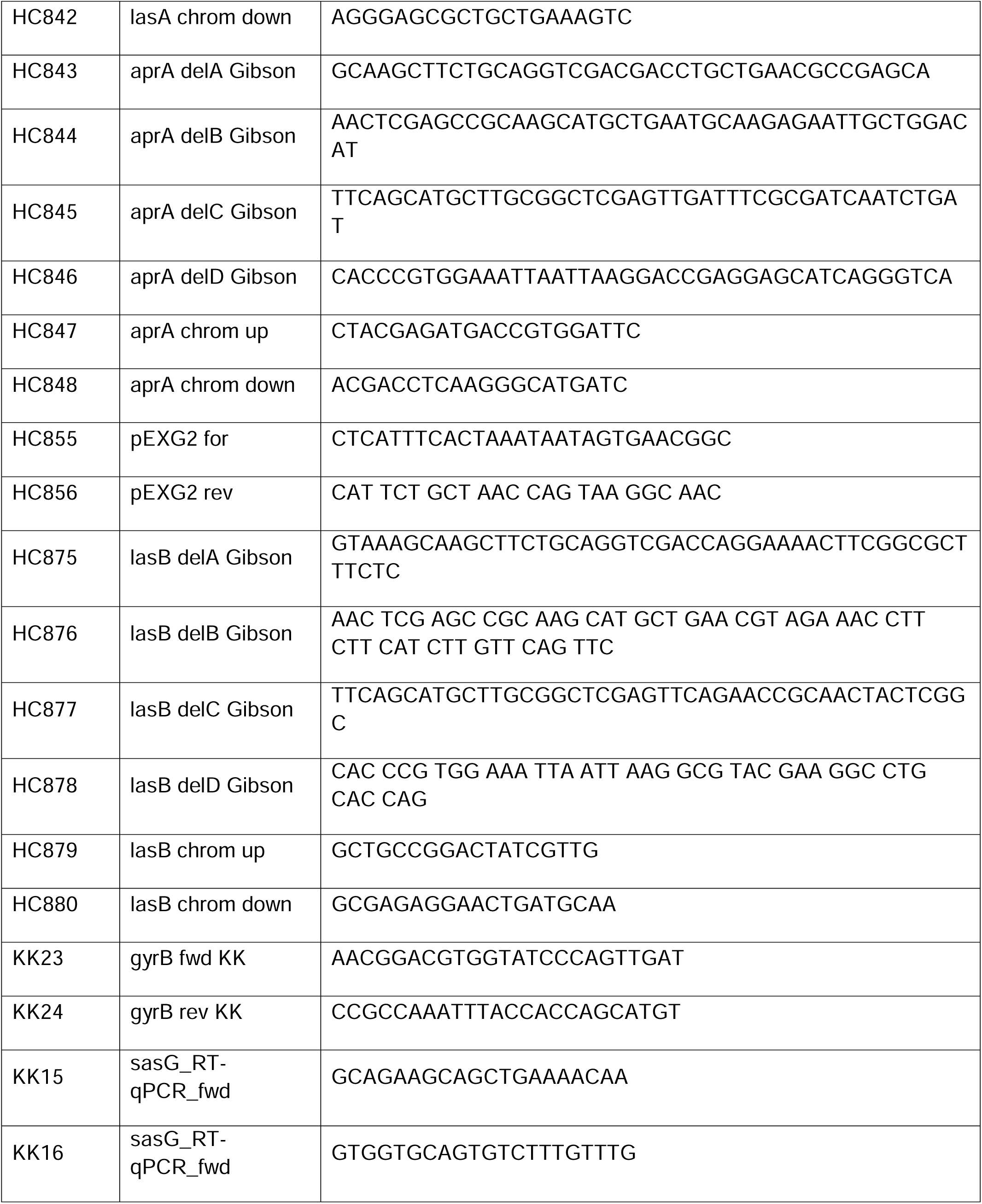
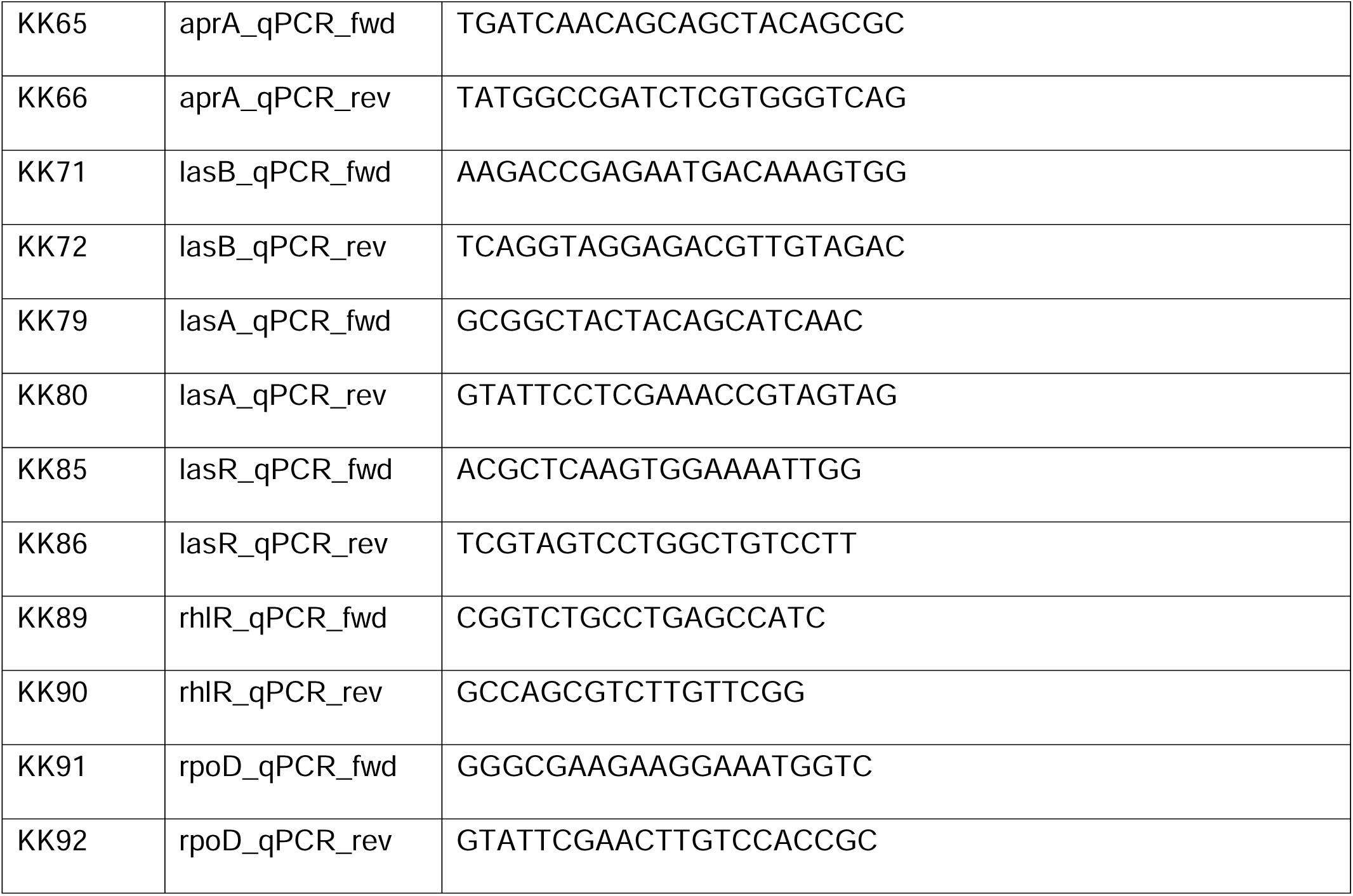
Primers.

### Construction of Fluorescent Reporter Strains

*S. aureus* MW2 *mgrA* and *sasG* mutant strains were fluorescently labelled by moving dsRed-expressing plasmid pHC48 [103] from RN4220 by phage transduction. Transductions were performed with phage 11 as described previously [104] and pHC48 was maintained with 10 µg/mL Cam. *P. aeruginosa* strain PAO1 was fluorescently labeled by moving GFP-expressing plasmid pMRP9-1 through electroporation and maintaining the plasmid with 100 µg/mL Amp[105].

### Construction of Protease Deletion Mutants in P. aeruginosa

Protease deletion mutants of *lasA, lasB, and aprA* were constructed in PAO1 through homologous recombination following conjugation as described previously[106]. The deletion constructs were generated in pEXG2 by Gibson assembly. Plasmid pEXG2 was purified from *E. coli* DC10B using the QIAprep Spin Miniprep kit (Qiagen). To generate the *lasA, lasB, and aprA* deletion plasmids, DNA fragments (∼700 bp in size) flanking the targeted regions for deletion and pEXG2 were amplified by PCR. The products were then purified with the QIAquick PCR Purification Kit (Qiagen), fused by a second amplification, and purified with the QIAquick Gel Extraction Kit (Qiagen). This PCR product and pEXG2 were digested with restriction enzymes, and ligated together to generate pHC207, pHC208, and pHC211. Plasmids were then electroporated into *E. coli* DC10B and plated on LB plates containing 20 µg/mL Gn to select for cells containing the plasmid. Single colonies were picked and patched on LB plates containing 20 µg/mL Gn. Presence of the plasmid with the flanking regions was then confirmed by PCR. The deletion plasmids were then purified from overnight cultures by miniprep and the insert was confirmed by sequencing.

Deletion plasmids were then moved from *E. coli* DC10B to *P. aeruginosa* PAO1 by triparental mating as described previously[106]. Single recombinants were selected for by plating on Vogel-Bonner minimal medium (VBMM) agar containing 160 µg/mL Gn and generate merodiploid strains[106]. Plasmids were resolved from merodiploid strains through counterselection by plating on VBMM containing 7.5% sucrose following overnight outgrowth in LB broth. Colonies were patched on VBMM agar containing 7.5% sucrose and screened for gene deletions by PCR, then deletions were confirmed by sequencing.

### Construction of an AprA expression strain in E. coli

To construct the *E. coli* BL21 strain expressing mature active AprA from *P. aeruginosa*, the *apr* operon was amplified by PCR from PAO1 genomic DNA using primers flanking upstream of *aprX* and downstream of *aprA*. All primers are listed in **Table 2**. The amplification products were PCR purified, and the plasmid pBAD18 was purified by miniprep. The purified *apr* operon insert and pBAD18 vector were digested with restriction enzymes and ligated together. The ligation product was then transformed into *E. coli* DH5α and plated on LB agar containing 100 µg/mL Amp for selection. Colonies were patched on LB agar containing 100 µg/mL Amp and screened by PCR for the presence of the plasmid with the insert. The plasmid was purified from overnight cultures and the insert was confirmed by sequencing. The plasmid was then transformed into *E. coli* BL21 by electroporation for protein expression.

### S. aureus Aggregation Assay

Cultures of *S. aureus* and *P. aeruginosa* (25mL) were grown overnight in TSB and LB, respectively, at 37°C with shaking at 200 rpm. One mL of *S. aureus* overnight culture was harvested by centrifugation and the supernatant was discarded. Cells were washed with 1 mL of phosphate buffered saline (PBS) and the centrifugation step was repeated, discarding the supernatant. *P. aeruginosa* cultures were centrifuged, the supernatants collected, and remaining cells removed with a 0.22 µm PVDF syringe filter (Nanopore). *P. aeruginosa* supernatants were then diluted in PBS to the appropriate concentration. *S. aureus* cells were resuspended in 1 mL of either *P. aeruginosa* cell-free supernatant or PBS. Tubes were allowed to sit for 1 hour at room temperature. Aggregation was assessed visually and quantified by optical density. Aggregation of *S. aureus* cells over 1 h results in sedimentation of the aggregates and clearing of the suspension. To quantify aggregation, 125µL of liquid was removed from the top of the suspension at timepoints 0 h and 1 h, and optical density at 600 nm was measured in a 96-well plate with a Tecan Infinite M200 plate reader. Results represent an average of three separate experiments with each performed in technical triplicate.

### Cell Wall Preparations

Following 1 h aggregation assays described above, *S. aureus* cell wall proteins were extracted as described previously[48]. Cells from aggregation assays were harvested by centrifugation, washed twice with PBS, and resuspended in 500 µL of protoplasting buffer (10mM Tris pH 8, 10mM MgSO_4_, 30% raffinose). Lysostaphin was added to the suspension and cells were incubated for 1 h at 37°C. Tubes were then centrifuged for 3 minutes a maximum speed and 500µL of supernatant was transferred to a new tube. Proteins were precipitated by addition of 125 µL of ice-cold trichloroacetic acid (TCA) and incubated on ice for 2 h. Precipitated proteins were centrifuged for 10 min at maximum speed and supernatant was discarded. The pellet was washed twice with 500 µL of ice cold 100% ethanol, centrifuging for 5 mins between washes and discarding supernatant, and inverted to dry. Pellets were resuspended in 36 µL 2X Laemelli SDS-PAGE Buffer (New England Biolabs), heated at 85°C, and loaded into 4-20% gradient acrylamide gel. Following SDS-PAGE, gels were stained with Coomassie blue stain and imaged.

### SasG Proteolytic Processing Assays and Cleavage Site Determination

*S. aureus* surface protein SasG was purified as described previously[48]. To assess SasG proteolysis, purified full-length SasG was diluted 10-fold in PBS to a concentration of 500 µg/mL. Then 2 µL of this SasG dilution was mixed with 18 µL of *P. aeruginosa* cell-free supernatant diluted in PBS to a final concentration of 1%. Reactions were incubated at room temperature for 10 minutes unless otherwise indicated. Reactions were quenched by adding 20µL of 2X Laemelli SDS-PAGE loading buffer (BioRad). Immediately following addition of loading buffer, 10µL was loaded on a 4-20% gradient gel. Following SDS-PAGE, gels were stained with Coomassie and imaged. SasG cleavage was quantified with ImageJ.

To determine the proteolytic cleavage site(s) in SasG, large-scale reactions were set up for each condition by mixing 30 µL SasG with 270 µL *P. aeruginosa* supernatant. Reactions were repeated as described above and 20 µL of reaction was loaded on a 4-20% gradient gel. Proteins were then transferred to a PVDF membrane using the Transblot Turbo Transfer System (BioRad) and the membrane was stained with Coomassie and dried. N-terminal sequencing was then carried out by Edman degradation using a Shimadzu PPSQ-53A Gradient Protein Sequencer at the Protein Facility at Iowa State University.

### RT-qPCR

For relative real-time quantitative PCR (RT-qPCR) quantification of *lasA, lasB,* and *aprA* expression, total RNA was isolated from *P. aeruginosa* using the Rneasy Mini Kit (Qiagen) according to the manufacturer’s instructions. Contaminating DNA was removed using Turbo DNA-free kit (Thermo Fisher). After DNase treatment, one step reverse transcription and real-time PCR amplification was performed on 100 ng of purified RNA using the iScript cDNA synthesis kit (Bio389 Rad). qPCR was performed by amplifying cDNA in 20 µL reaction volumes with iTaq Universal SYBR Green Supermix (Bio-Rad) in the CFX96 Touch Real-Time PCR System (Bio393 Rad) under the following conditions: 3 min at 95°C, 40 cycles of 10 s at 95°C and 30 s at 60°C, followed by a dissociation curve. No template and no reverse transcription controls were performed in parallel. Primers used for the amplification of *aprA, lasB*, *lasA* and *rpoD* are described in **Table 2**.

Results reflect three independent experiments performed in triplicate. Relative expression was normalized to *rpoD* via the Pfaffl method.

### Assessment of Antimicrobial Resistance

The MICs of vancomycin (Van) and ciprofloxacin (Cip) were determined for each *S. aureus* strain by the standard broth microdilution method according to CLSI guidelines[107, 108]. The MICs were estimated accordingly: Van = 1.0 µg/mL and Cip = 0.5 µg/mL. MIC did not vary among mutant strains of *S. aureus* MW2 and estimations were consistent with the EUCAST predicted MICs. To assess changes in antimicrobial susceptibility, aggregation assays were performed as described above. Following aggregation for 1hr, tubes were centrifuged and supernatant discarded. Cells were resuspended in 1 mL of CAMHB and mixed gently by pipetting. Antibiotic-supplemented media was prepared by diluting vancomycin or ciprofloxacin in CAMHB to final concentrations of 0, 1, 2, and 4 µg/mL. Antibiotic-supplemented CAMHB was inoculated with ∼5×10^6^ CFU of *S. aureus* to a final volume of 200 µL in a 96-well plate, and each plate included both sterility and growth controls. An initial time point (t = 0 h) was taken by plating for CFUs on Cation-Adjusted Mueller-Hinton Agar (CAMHA). Plates were incubated at 37°C, with timepoints taken at t = 1, 3, and 5 h and CFUs enumerated. Growth was also assessed by measuring OD_600_ at each timepoint in addition to t= 10 and 18 h (**Supplementary Figure 2)**. Results represent four separate experiments, and each condition was performed in triplicate.

### In vitro Lubbock Chronic Wound Biofilm Model

In this study, we adapted the Lubbock Chronic Wound Biofilm Model developed by Sun et al.[81]. Wound-like Media (WLM; 50% Bolton broth, 45% heparinized bovine plasma, 5% laked horse blood) was aliquoted (3mL) into sterile glass test tubes. Overnight cultures (5 mL) of *S. aureus* and *P. aeruginosa* were normalized to an OD_600_ of 0.125 in Bolton broth (BB). This suspension was subsequently diluted 1:10 into 900 µL BB so that 10µL of each culture was normalized to 5×10^5^ CFU. LCWBM preparation followed by inoculating WLM with 10 µL of diluted *S. aureus* and *P. aeruginosa* as monocultures and cocultures. A sterile pipette tip (20µL Rainin SoftFit-L Tips; Thermo Fisher Scientific) was ejected into the test tube during inoculation. Biofilms were cultured at 37°C with shaking and harvested after 24 h of incubation. Biofilms were harvested from the glass tubes, imaged, and the pipette tip was removed. Each biofilm was washed three times with 500 µL sterile PBS and transferred to a new sterile plate, imaged, and excess medium was removed. Biofilms were transferred to sterile pre-weighed tubes containing four steel homogenization beads and 500µL sterile PBS. Tubes were bead-beat for 90 s at three 30 s intervals, with tubes placed on ice for 30 s between each bead-beating. Tubes were vortexed for 1 min, and CFU/mg was determined by serial dilution and selectively plating for *S. aureus* on Mannitol Salt Agar (MSA) and *P. aeruginosa* on *Pseudomonas* Isolation Agar (PIA). Results represent three separate experiments with each condition performed in triplicate.

### Confocal Laser Scanning Microscopy and Image Analysis

Lubbock Chronic Wound Biofilms were additionally analyzed by confocal laser scanning microscopy (CLSM) using the Olympus FV1000-IX81 Microscope at the University of Colorado Anschutz Medical Campus Advanced Light Microscopy Core. Biofilms were cultured as described above using dsRed-expressing *S. aureus* strains (pHC48) and GFP-expressing *P. aeruginosa* (pMRP9-1). Following removal of the pipette tip scaffold, harvested biofilms were placed on glass slides, fixed with 10% formalin, and coverslips were placed carefully to cover the biofilm. Detection of dsRed-expressing *S. aureus* cells was performed using excitation/emission wavelengths of 587/610 nm. Detection of GFP-expressing *P. aeruginosa* was performed by using excitation/emission wavelengths of 488/509 nm. Images were acquired using 20×, 60× water-immersion, and 100× oil-immersion objectives. Data were stored as 1024-by 1024-pixel slices in stacks of 20 images. Three biofilms were imaged for each condition and results reflect the most representative images of each condition.

### Murine Model of Polymicrobial Chronic Wound Infections

All animals are housed and maintained at the University of Colorado Anschutz Medical Campus Animal Care Facility accredited by the Association for Assessment and Accreditation of Laboratory Care International (AAALAC). All animal studies described herein were performed in accordance with best practices outlined by the Office of Laboratory Animal Resources (OLAR) and Institutional Animal Care and Use Committee (IACUC) at the University of Colorado (protocol #00987). *S. aureus* MW2 strains and *P. aeruginosa* strain PAO1 were grown overnight at 37°C with shaking in 5mL of TSB and LB, respectively. Overnight cultures were diluted 1:100 into flasks containing 35 mL of TSB and LB and subcultured at 37°C with shaking to an OD_600_ of 0.5. Subcultured bacteria were then pelleted, and resuspended in sterile saline so that all strains were normalized to 5×10^5^ CFU/10 µL. One mL of each strain was aliquoted into an Eppendorf tube and kept on ice throughout the experiment.

A murine model of polymicrobial chronic wound infection was used to assess persistence and infection dynamics between *S. aureus* and *P. aeruginosa.* C57BL/6 female mice (Jackson Laboratories) arrived to the animal facility at 7-weeks of age. Mice were allowed to acclimate to the BSL-2 level animal housing facility at the University of Colorado Anschutz Medical Campus for at least seven days prior to their inclusion in this study’s *in vivo* infection model. One day prior to infection, mice were anesthetized (2-3% isoflurane; inhalation) and fur on the dorsal surface was carefully shaved. Nair was applied to remove any remove any remaining fur and completely expose the skin. On day 0, mice were anesthetized and the shaved skin surface was sterilized with an isopropyl alcohol swab and povidone iodine prep pad (PDI Healthcare). Bupivacaine hydrochloride was used as a local anesthetic for the area to be wounded and was injected subcutaneously at a dosage of 1-2 mg/kg. Buprenorphine was used as an analgesic and was injected subcutaneously at a dosage of 0.01-0.2mg/kg. A 6mm biopsy punch was used with dissection scissors and forceps to excise a circular section of skin and generate a wound 6 mm in diameter. Following wounding, each mouse was inoculated by pipetting a final volume of 10 µL of bacterial inoculum or sterile saline (vehicle control) directly onto the wound. Single infections were inoculated with 10µL (5 x 10^5^ CFU) of *S. aureus* or *P. aeruginosa.* Co-infections were inoculated with 5 µL (2.5 x 10^5^ CFU) of both *S. aureus* and *P. aeruginosa*. Following infection, wounds were covered with the transparent dressing Tegaderm followed by two bandages.

The infection time course spanned nine days, Tegaderm was removed on day 2, and bandages were replaced daily. Clinical severity was assessed by measuring body weight changes. Lesions were imaged to assess wound severity and healing progression and was analyzed with ImageJ software (National Institutes of Health). On day 9, animals were euthanized by CO2 inhalation followed by cervical dislocation. Wound tissue was excised and placed in a pre-weighed 2mL vial with 0.5mL of 1XPBS and 1.0mm zirconia/silica beads. The excised tissue was homogenized by bead-beating for 90 s at three 30 s intervals, with tubes placed on ice for 30 s between each bead-beating. Tubes were vortexed for 1 min, and CFU/mg was determined by serial dilution and selectively plating for *S. aureus* on Mannitol Salt Agar (MSA) and *P. aeruginosa* on *Pseudomonas* Isolation Agar (PIA). Results represent three separate experiments with five mice per condition.

## Resource availability

Further inquiries and information on reagents and resources should be directed to (and will be fulfilled by) the lead contact, Alexander R. Horswill.

## Acknowledgements

We thank the members of the Horswill and Doran labs at the University of Colorado Medical School, for their critical evaluation of this work.

A.R.H is funded by NIH award AI083211 and the Department of Veteran’s Affairs award BX002711.

## Author Contributions

Conceptualization, K.K., M.B., C.J., M.S., A.R.H.; Methodology, K.K., M.B., C.J., H.C., K.M., A.R.H.; Investigation, K.K., C.J.; Writing-Original Draft, K.K., Writing-Review and Editing, K.K., M.B., C.J., H.C., M.S., A.R.H.; Funding Acquisition, A.R.H.; Supervision M.S., A.R.H.

## Supplementary Figures

**Supplementary Figure 1.**
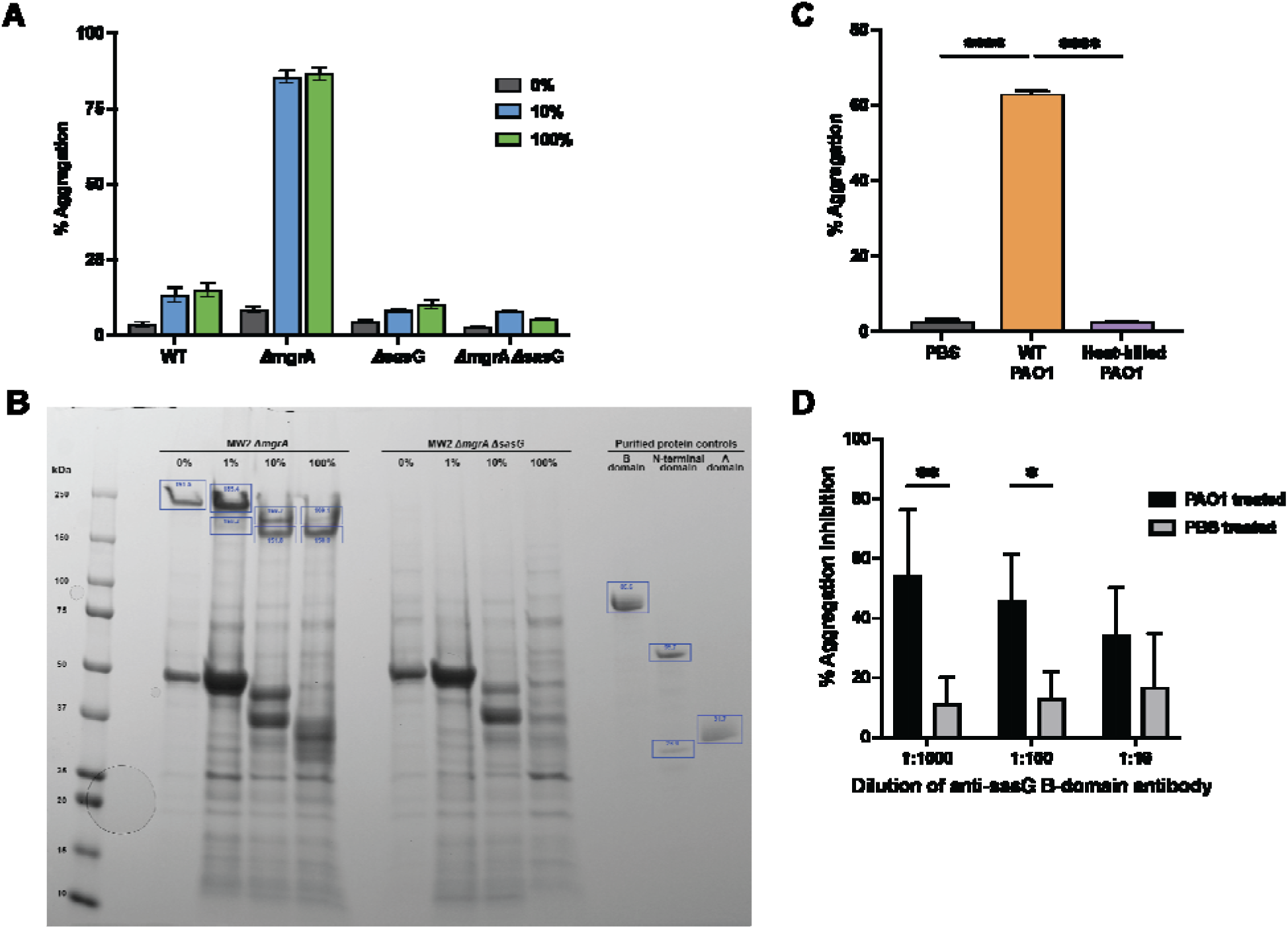
The regulator MgrA represses *sasG* under laboratory conditions. (A) Aggregation of MRSA strain USA400 MW2 WT, Δ*mgrA,* and Δ*sasG* strains treated with 0-100% WT PAO1 supernatant. (B) Quantification of Coomassie stained SDS-PAGE gel showing cell wall protein extractions from Figure 1D following aggregation. (C) MRSA Δ*mgrA* mutant aggregation treated with 10% heat treated and WT PAO1 supernatant. (D) Blocking of *S. aureus* aggregation by SasG B domain antibodies. Results represent an average of three independent experiments performed in triplicate ± SEM (n=6). Statistical significance was determined by one-way ANOVA with Bonferroni multiple comparisons test (****p<0.0001).

**Supplementary Figure 2.**
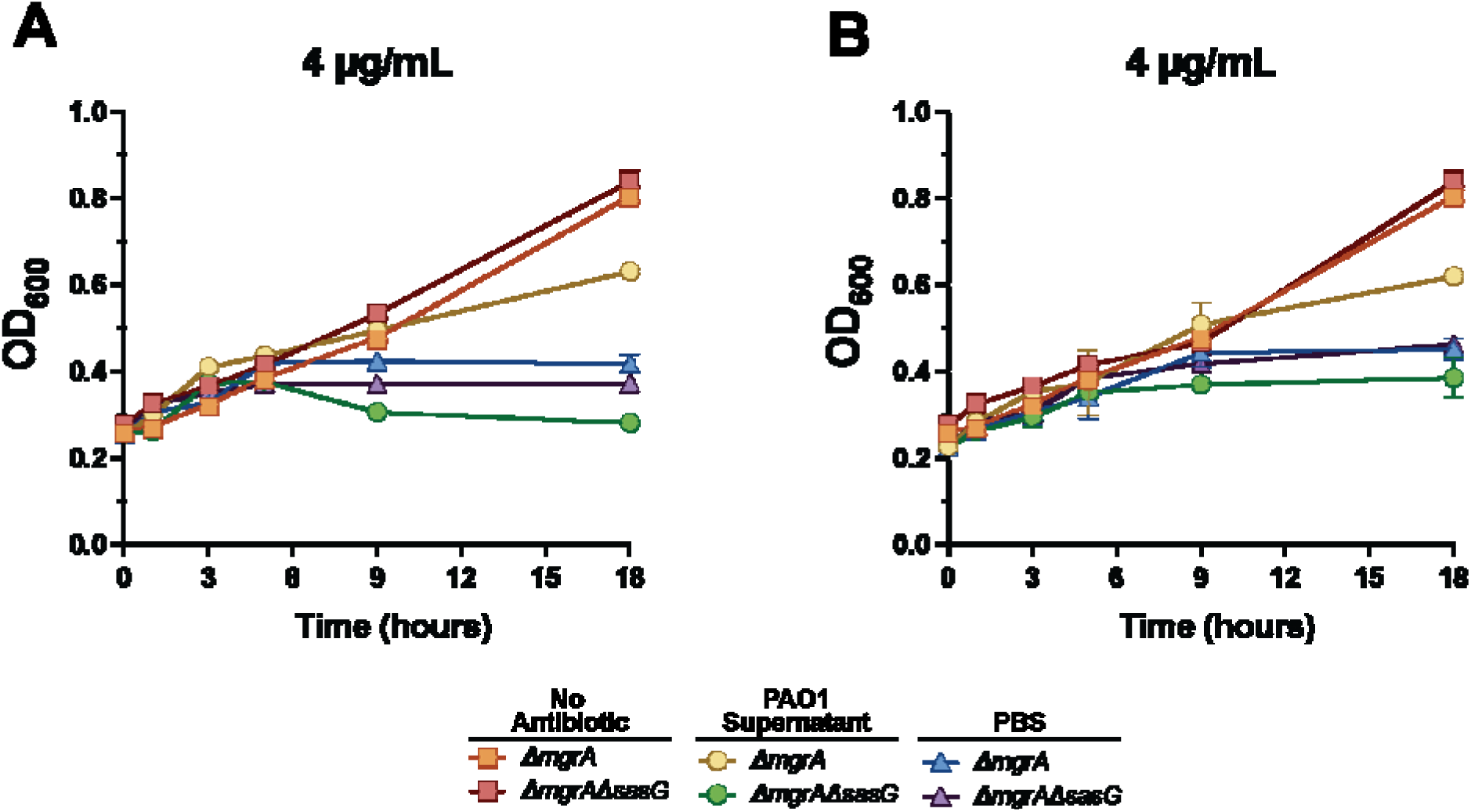
SasG-dependent MRSA aggregates are more tolerant to antimicrobials. OD600 showing growth of MRSA over 18 hours with treatment of (A) Cip or (B) Vn.

**Supplementary Figure 3.**
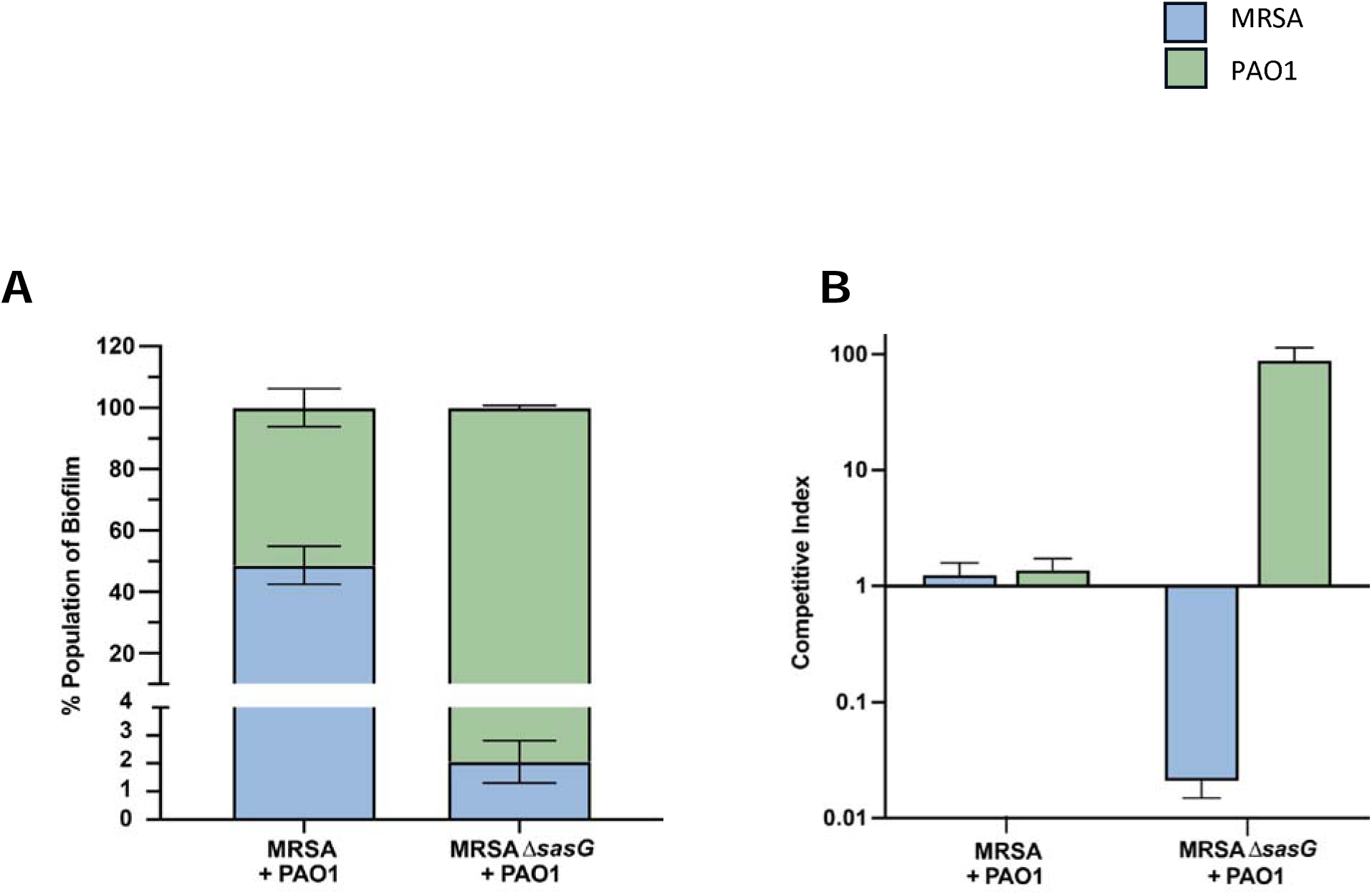
Quantification of population dynamics in Lubbock polymicrobial biofilms. (A) Population makeup of Lubbock polymicrobial biofilms. (B) Competitive Index between MRSA and PAO1 in polymicrobial biofilms. All data analyzed with COMSTAT. Results represent an average of three independent experiments performed in triplicate ± SEM (n=9).

